# Genomic characterization of four novel bacteriophages infecting the clinical pathogen *Klebsiella pneumoniae*

**DOI:** 10.1101/2021.03.02.433402

**Authors:** Boris Estrada Bonilla, Ana Rita Costa, Teunke van Rossum, Stefan Hagedoorn, Hielke Walinga, Minfeng Xiao, Wenchen Song, Pieter-Jan Haas, Franklin L. Nobrega, Stan J.J. Brouns

## Abstract

Bacteriophages are an invaluable source of novel genetic diversity. Sequencing of phage genomes can reveal new proteins with potential uses as biotechnological and medical tools, and help unravel the diversity of biological mechanisms employed by phages to take over the host during viral infection. Aiming to expand the available collection of phage genomes, we have isolated, sequenced, and assembled the genome sequences of four phages that infect the clinical pathogen *Klebsiella pneumoniae:* vB_KpnP_FBKp16, vB_KpnP_FBKp27, vB_KpnM_FBKp34, and Jumbo phage vB_KpnM_FBKp24. The four phages show very low (0-13%) identity to genomic phage sequences deposited in the Genbank database. Three of the four phages encode tRNAs and have a GC content very dissimilar to that of the host. Importantly, the genome sequences of the phages reveal potentially novel DNA packaging mechanisms as well as distinct clades of tubulin spindle and nucleus shell proteins that some phages use to compartmentalize viral replication. Overall, this study contributes to uncovering previously unknown virus diversity, and provides novel candidates for phage therapy applications against antibiotic-resistant *K. pneumoniae* infections.

## 1. Introduction

Bacteriophages or phages are ubiquitous viruses of prokaryotes that exert an enormous influence over the microbial biosphere, playing a critical role in the nutrient and energy cycles,^1–3^ in the evolution of bacterial pathogens,^4^ and in shaping gut microbial communities.^5^ Phages have also contributed immensely to the field of molecular biology, having been at the core of the discovery of central features such as DNA as the genetic material,^6^the triplet genetic code,^7^messenger RNA,^8^ restriction enzymes,^9^and recombinant DNA.^10^ Phages and their interactions with prokaryotic hosts led also to the evolution of CRISPR-Cas and development of programmable genome editing tools, one of the most revolutionary tools in biology that enables tailored engineering of genomic sequences in a range of species including humans.^11^There is also a rekindled interest in the therapeutic use of phages – phage therapy – to control bacterial pathogens, as a consequence of the alarming rise of antibiotic resistant infections observed in recent years.^12–14^ The study of phages and their genomes is therefore inherently valuable to advance our understanding in a diversity of fields including molecular biology, ecology, evolution, bacterial pathogenesis, biotechnology and health. Understanding phage genomes will certainly create opportunities to translate novel phage proteins and phages themselves into potent biotechnological^15^ and medical tools.^16^ Here we isolated and sequenced the genomes of four novel phages infecting *Klebsiella pneumoniae*, an increasingly relevant pathogen identified by the World Health Organization as priority for the development of new antibiotics.^17^ These phages have little to no sequence similarity to known phages, but a series of genomics and phylogenetic analysis revealed interesting features that could aid the expansion of our understanding of the hidden genetic treasures in phage biology.

## 2. Materials and methods

### 2.1. Bacteriophage isolation

Four clinical isolates of *K. pneumoniae* isolated at the University Medical Centre Utrecht (UMCU) were used for phage isolation: K6310 (blood culture from a 77 year-old patient with obstructive cholangitis due to disseminated pancreatic carcinoma), K6592 (infected total hip prosthesis from a 74 year-old patient), L923 (blood culture from 67 year-old kidney transplant patient with an urinary tract infection and sepsis) and K6453 (cerebrospinal fluid taken post-mortem from a healthy 57 year-old woman with unexplained sudden out of hospital cardiac arrest). As phage source, approximately 5 L of sewage water were sequentially filtered with coffee filters, membrane filters (0.45 and 0.2 µm PES), and 10x concentrated using a tangential flow cassette (100 kDa PES Vivaflow 200, Sartorius, Germany). Approximately 5 mL of the concentrated virome were added to 20 mL of Lysogeny Broth (LB), inoculated with 100 µL of each of the overnight grown *K. pneumoniae* strains, and incubated overnight at 37 °C, 180 rpm. Samples were centrifuged at 16,000 × *g* for 5 min and filter-sterilized (0.2 µm PES). The phage-containing supernatant was serially diluted in SM buffer (100 mM NaCl, 8 mM MgSO_4_, 50 mM Tris-HCl pH 7.5) and spotted on double layer agar (DLA) plates of the isolation strains for the detection of phages. Single plaques with distinct morphologies were picked with sterile toothpicks and spread with sterile paper strips into fresh bacterial lawns. The procedure was repeated as needed to obtain a consistent plaque morphology. Phages from purified plaques were then produced in liquid media with their respective host, centrifuged, filter-sterilized and stored as phage lysates (>10^8^ pfu/ml) at 4°C, and at -80°C with 50 % (v/v) glycerol.

### 2.2. Transmission electron microscopy

One mL of each phage lysate at >10^9^ pfu/mL was sedimented at 21,000 × *g* for 1 h, washed and re-suspended in 1 mL of MilliQ water. Phages (3.5 µL) were deposited and incubated for 1 min on TEM grids (Carbon Type-B 400 mesh, TED PELLA). Grids were washed thrice with 40 µL of MilliQ water and stained with 3.5 µL of 2% (w/v) uranyl acetate (pH 4.0) for 30 seconds. Grids were examined using a JEM-1400 plus (JEOL) TEM. The capsid diameter and the tail length and width of 10 phage particles were measured using EMMENU v4.0.9.8.7 (Tietz Video & Image Processing Systems GmbH, Gauting, Germany) and used to calculate the average dimensions of each phage.

### 2.3. Bacteriophage genome sequencing

Phage DNA was extracted using phenol-chloroform as previously described.^18^ DNA was sequenced by the BGI Group (Shenzhen, China) using the BGI MGISEQ-2000 platform. Quality control of the raw data was performed using FastP^19^ and Soapnuke,^20^ and the reads were trimmed and processed using Seqtk.^21^The filtered reads were assembled into the final genomes with SPAdes.^22^

### 2.4. Bacteriophage genome annotation and comparative genomics

Phage genomes were automatically annotated using the RAST server v2.0.^23^ Additional putative functions were assigned to coding sequences (CDS) by BlastP v.2.10.0^24^ and Hmmer v3.3.1.^25^ tRNAs were predicted with tRNAscan-SE v2.0^26^. Rho-independent terminators and promoters were identified with ARNold^27^ and PhagePromoter v0.1.0^28^, respectively. Genomic comparisons were performed using BlastN v.2.10.0^24^. Schematics of phage genomes were built with CGView Server^29^.

### 2.4. Evolutionary analysis of phage proteins

The genome packaging strategy of the phages was predicted by phylogenetic analysis of the large terminase subunit as previously described.^30^ Evolutionary relationships of phage tubulin spindle and nucleus shell proteins were investigated by building phylogenetic trees with proteins found by psi-Blast (with five iterations) and Hmmer to be homologous to the tubulin spindle (**Supplementary Table S1**) and nucleus shell (**Supplementary Table S2**) proteins of *Pseudomonas* phage 201phi2-1, with an e-value equal or less than 1e-5. For all trees, proteins were aligned using MAFFT v7.308 with default settings, and the trees built using RAxML 7.2.8 with bootstrapping set to 100. A consensus tree was obtained using Consensus Tree Builder in Geneious v9.1.8.

### 2.5. Codon usage analysis

Codon usage of the bacteriophages and the *K. pneumoniae* HS11286 reference genome (GenBank RefSeq: NC_016845.1) was analyzed with Cusp from EMBOSS.^31^

## 3. Results and discussion

### 3.1. General morphological and genomic features

We have isolated four phages infecting *K. pneumoniae* from sewage samples: vB_KpP_FBKp16 (ϕKp16), vB_KpP_FBKp27 (ϕKp27), vB_KpM_FBKp34 (ϕKp34) and vB_KpM_FBKp24 (ϕKp24). The four phages have a tail and therefore belong to the *Caudovirales* order of phages with double stranded DNA. Phages ϕKp16 (**Fig. 1A**) and ϕKp27 (**Fig. 1B**) have short tails and encode an RNA polymerase (**Supplementary Tables S3 and S4**), features that classify these phages in the *Autographiviridae* family.^32^ Phage ϕKp34 (**Fig. 1C**) has a long contractile tail and a small baseplate with tail spikes and no tail fibers, a distinctive feature of *Ackermannviridae*.^33^ Phage ϕKp24 (**Fig. 1D**) also has a long contractile tail, but with a complex tail fiber structure at the baseplate, and a capsid that is 1.5 times larger than that of ϕKp34. These morphological features and the large ≈307 kb genome (**Table 1**) indicate that ϕKp24 is a Jumbo *Myoviridae*.

**Table 1.**
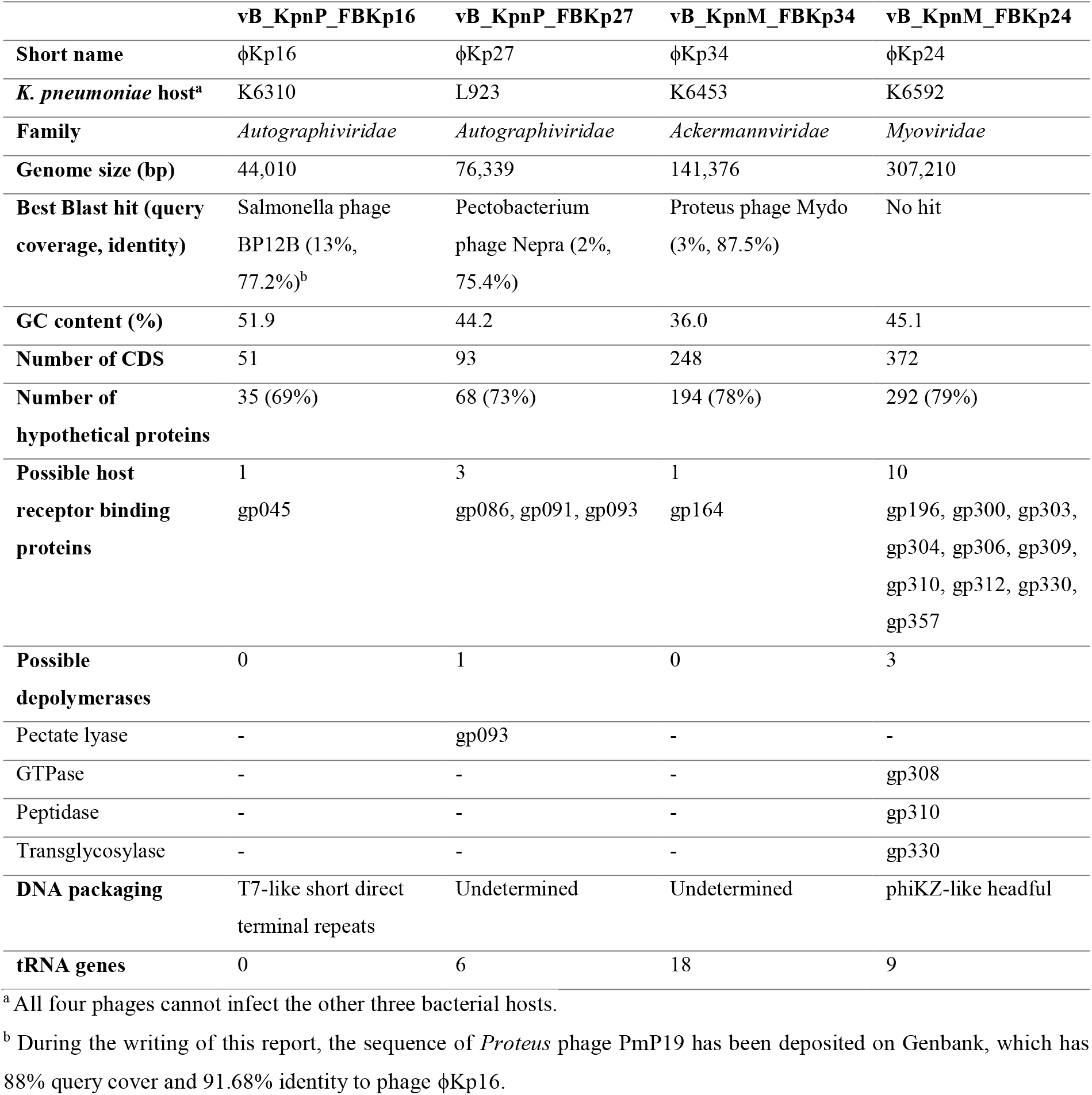
Morphological and genomic features of the bacteriophages isolated in this work.

**Figure 1.**
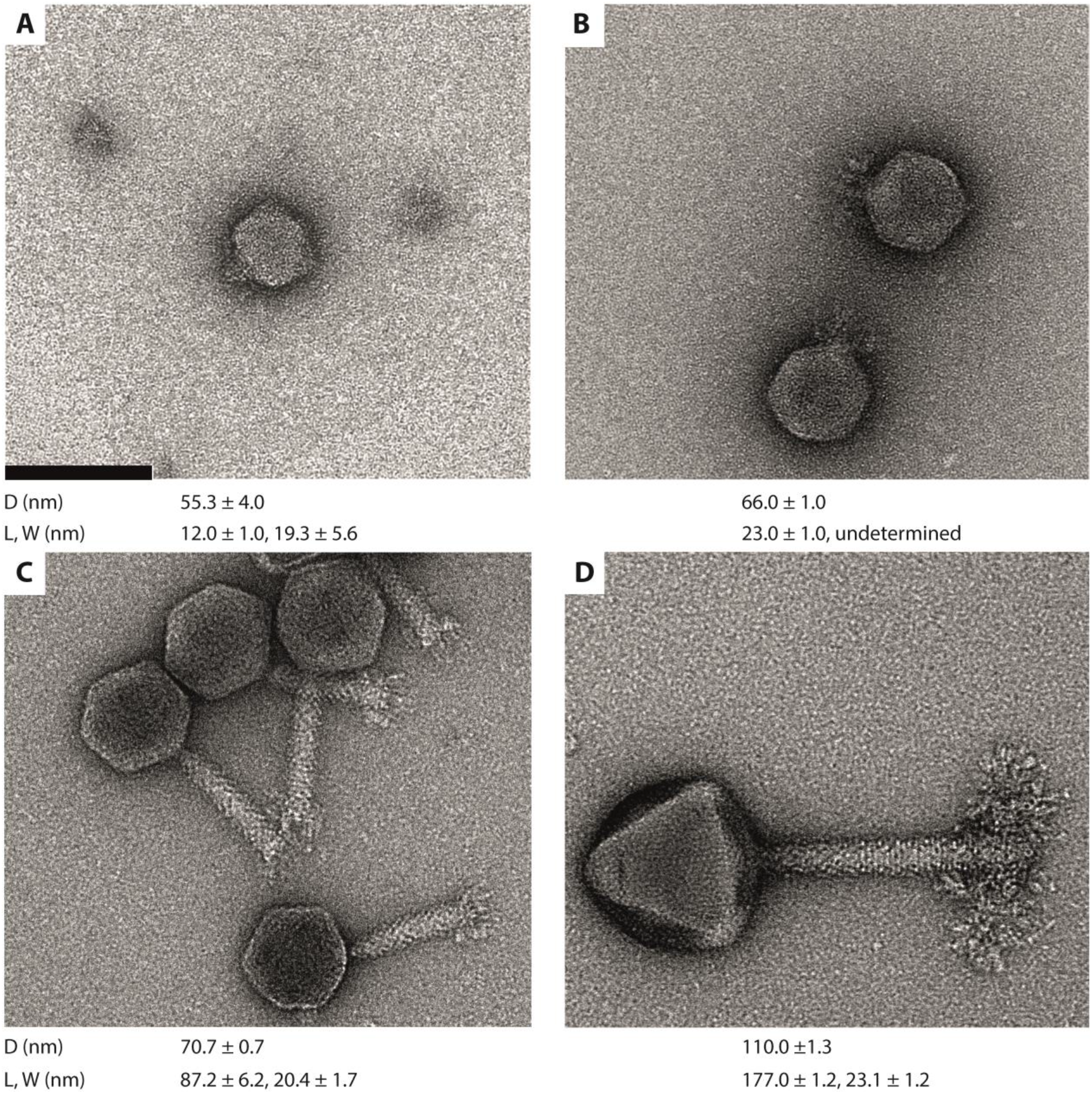
Transmission electron microscopy images of four newly isolated *Klebsiella pneumoniae* bacteriophages. (A) *Autographiviridae* ϕKp16, (B) *Autographiviridae* ϕKp27, (C) *Ackermannviridae* ϕKp34, and D) *Myoviridae* ϕKp24. Bacteriophages were negatively stained with 2% uracyl acetate. The diameter (D) of the capsid, and the length (L) and width (W) of the tail are given in nm bellow each phage as the average dimensions of 10 phage particles. Bar: 100 nm. All micrographs are taken at 200,000x magnification.

The four phages share a very low sequence similarity to each other and to phage genome sequences deposited in Genbank (**Table 1**). The genome of phage ϕKp24 is of particular highlight since no similarity was found to any genome sequence in Genbank, underlining its novelty. The genomes of phages are often organized in clusters of functionally related genes. While this analysis is made difficult due to the high number of hypothetical proteins with unassigned function (69-79%, **Table 1**), it is still possible to observe functional gene clustering. All genes but one in phage ϕKp16 are oriented in the same direction and organized in functional groups, especially evidenced by DNA replication and repair, and structural component genes (**Fig. 2A, Supplementary Table S3**). Genes in phage ϕKp27 are organized in clusters of different orientation that group genes of related functions (**Fig. 2B, Supplementary Table S4**). Cluster A groups genes involved in DNA metabolism, while Cluster B groups genes for transcription (an RNA polymerase) and a first set of genes for structural components related to capsid and tail tape measure proteins. Cluster C contains genes involved in regulation and six tRNAs, while Cluster D has only one gene with function identified for DNA packaging. Cluster E groups a second set of genes for transcription (a second RNA polymerase) as well as genes involved in DNA metabolism and DNA replication and repair. Finally, cluster F groups genes for a second set of structural components related to tail and host binding proteins, as well as genes for cell lysis. Genes in phage ϕKp34 are also organized in clusters of opposing orientation grouping genes of related functions, although genes for similar functions appear in more than one cluster (**Fig. 2C, Supplementary Table S5**). Cluster A groups genes related to DNA methylation, DNA metabolism and DNA replication and repair. Cluster B also groups genes for DNA metabolism and DNA replication and repair, as well as multiple genes seemingly related to bacterial metabolism (**Supplementary Table S5**). Cluster C groups all genes identified as structural components, as well as genes involved in DNA packaging, DNA integration, DNA metabolism and tRNAs, and Cluster D groups most genes related to DNA replication and repair, as well as genes for DNA metabolism, DNA methylation, DNA recombination, and cell lysis. Genes in phage ϕKp24 are mostly oriented in the same direction and in some cases the predominant gene orientation is reversed by individual or small groups of genes in the opposite orientation (**Fig. 2D, Supplementary Table S6**). The large genome size and the high percentage (79%) of proteins with unassigned functions in phage ϕKp24 make it difficult to define functional gene groups. Still, it is possible to identify three major groups of genes coding for structural components, as well as small groups of genes involved in DNA metabolism, transcription and DNA replication and repair, evidencing the functional clustering of genes commonly observed in phages.

**Figure 2.**
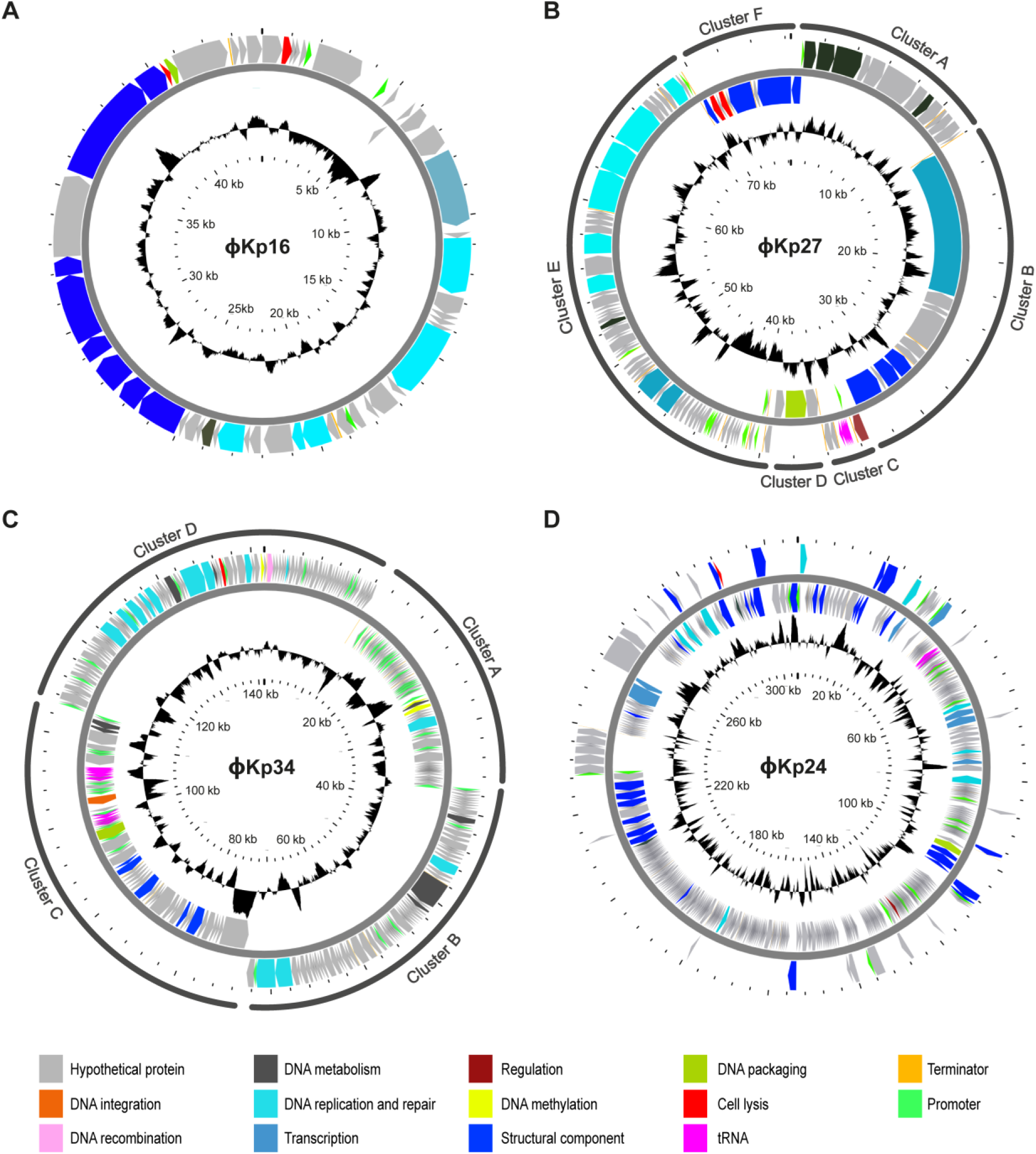
Circular genome maps of the four newly isolated *Klebsiella pneumoniae* bacteriophages. (A) Phage ϕKp16, (B) Phage ϕKp27, (C) Phage ϕKp34, and (D) Phage ϕKp24. ORFs are colored according to predicted function as shown in the key. Clusters depict clear gene operons located in the same strand. Clusters are not shown for (A) and (D) since most genes are located in the same strand. GC content is shown in black landscape with GC content higher or lower than 50% pointing to outer and inner circles, respectively. Maps were generated using CGView Server.^29^

Phages ϕKp27, ϕKp34, and ϕKp24 encode 6, 18 and 9 tRNA genes, respectively (**Table 1**). As of yet, there is no clear explanation for the presence of tRNA genes in phage genomes. ^34–36^ A number of studies have proposed that tRNA-containing phages have a codon bias that diverges from that of the bacterial host, therefore using the tRNAs to compensate for a metabolic difference.^37,38^ However, other studies have shown that this is not an universal observation. Here, we observe that less than half of the tRNAs encoded in phages ϕKp27 (3 of 6), ϕKp34 (7 of 18), and ϕKp24 (4 of 9) associate with codons that are more used in the phage than in the bacterial host (**Fig. 3, Supplementary Table S7**), suggesting that codon bias is not the (only) explanation for the presence of tRNAs in phages. It has also been suggested that tRNAs in phages may be beneficial to overcome the codon bias of different hosts,^39^but this is difficult to assess since it is virtually impossible to determine the full range of species and strains that a phage can infect. Interestingly, 67% (18 of 27) of the codons more highly expressed by the phages are shared by at least two phages, with 44% (12 of 27) being shared by the three. It will be interesting to explore common features of codon usage among phages of a certain species, rather than the similarity of codon usage between phage and host, as a feature to help predict the host in the future. Of note is also the presence of a suppressor tRNA (tRNA-Sup-TTA, **Supplementary Table S6**) in ϕKp24. Suppressor tRNAs arise when a mutation changes the tRNA anticodon, allowing it to recognize a stop codon and, instead of terminating, insert an amino acid at that position in the polypeptide chain.^40^ By doing so, suppressor tRNAs can give rise to abnormally long proteins and produce metabolic changes.^41^ In phages, suppressor tRNAs have been shown to alleviate nonsense mutations (formation of a non-functional protein due to the premature appearance of a terminator codon in mRNA) that sometimes appear due to the rapid mutation rate of phages.^42^ Whether the suppressor tRNA of phage ϕKp24 serves a similar or different (e.g. interfere with host protein expression for host takeover) purpose, requires further investigation.

**Figure 3.**
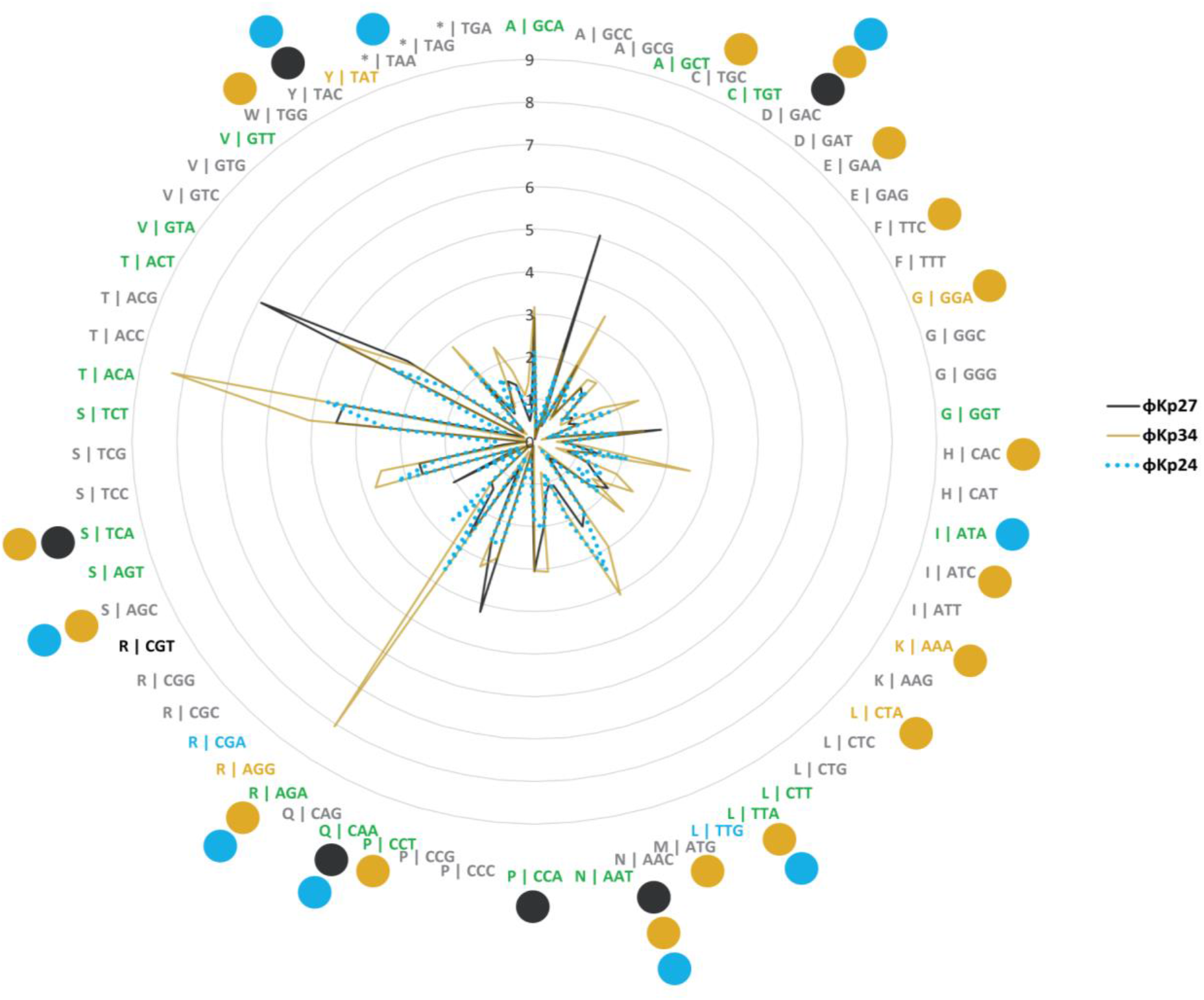
Codon usage by *Klebsiella pneumoniae* phages ϕKp27, ϕKp24 and ϕKp34 as compared to the codon usage of *K. pneumoniae* HS11286. Codon usage is represented as the fraction between the frequency of codon usage in the phage divided by that of the bacteria. Codons are represented as X |YYY, in which X is the amino acid codified by codon YYY. Codons expressed at least 2-fold higher in phages are colored black if overexpressed only in ϕKp27, mustard if only in ϕKp34, blue if only in ϕKp24, and green if overexpressed in at least two of the three phages. Colored circles indicate codons for which the phages encode a tRNA.

The GC content of the phages (36.0-51.9%) is lower than the median GC content of *K. pneumoniae* (57.2%), a feature that is particularly prominent for phage ϕKp34 (36.0%) (**Table 1**). These results corroborate previous studies that show the GC content of phage genomes to accurately (>95%) predict the host associated with a phage at the phyla level but not at lower taxonomic levels.^43^ In fact, the divergence in GC content between phages and their bacterial hosts has been previously observed for phages infecting different species.^44–47^ Curiously, phage ϕKp34 has the lowest GC content and encodes for the largest number of tRNA genes, while phage ϕKp16 has the GC content closest to its host and encodes no tRNA genes, suggesting a connection worth exploring in future work.

### 3.2. Bacteriophage ϕKp16 has internal virion proteins

Phage ϕKp16 encodes two proteins annotated as putative internal virion proteins (gp042 and gp044), i.e. proteins that are encapsidated with the phage genome in the phage capsid during phage assembly inside the cell. In particular, gp044 holds similarity to gp37 of Enterobacteria phage SP6 (99% query cover, 75.71% identity). This protein is a homologue of protein gp16 of Enterobacteria phage T7, which forms part of the ejectosome complex that degrades the bacterial cell wall prior to DNA ejection, by forming an inner pore in the inner membrane to allow entry of the phage DNA into the cell.^48^ Phage ϕKp16 most likely uses a similar mechanism in which proteins encapsidated with the genome are ejected to form a transmembrane channel through which the phage genome can cross to enter the cell cytoplasm. The proteins in ϕKp16 are however distantly related to those of phage T7 and even SP6, suggesting a possible variant mode of channel formation.

### 3.3. Bacteriophage ϕKp27 has a potentially novel genome packaging mechanism

Genome packaging is a critical step in the assembly of *Caudovirales* phages and is carried out by a protein known as the large terminase.^49^ Using a phylogenetics approach^30^ with the large terminase subunits of our phages and phages with well-characterized packaging mechanisms, we could infer the packaging mechanisms used by our phages (**Fig. 4A, Table 1**). The terminases of phages ϕKp16 and ϕKp24 clustered within groups of known packaging mechanisms, T7-like short direct terminal repeats and phiKZ-like headful packaging, respectively (**Fig. 4A**). However, the terminases of phages ϕKp27 and ϕKp34 formed their own clades, suggesting packaging mechanisms distinct from those currently known. Of these, the terminase of ϕKp27 seems to be the most distinctive, considering its distancing to all clades. A Blastp analysis revealed that the large terminase of ϕKp27 is highly similar (99% query cover, >70% identity) to the large terminase of N4-like phages.^50,51^The crystal structure of both the large and small terminase subunits of N4 have been resolved,^49^ but the mechanism of genome packaging is yet to be described and may reveal a mechanism different from those known so far, and which seems to be common to a number of phages.

**Figure 4.**
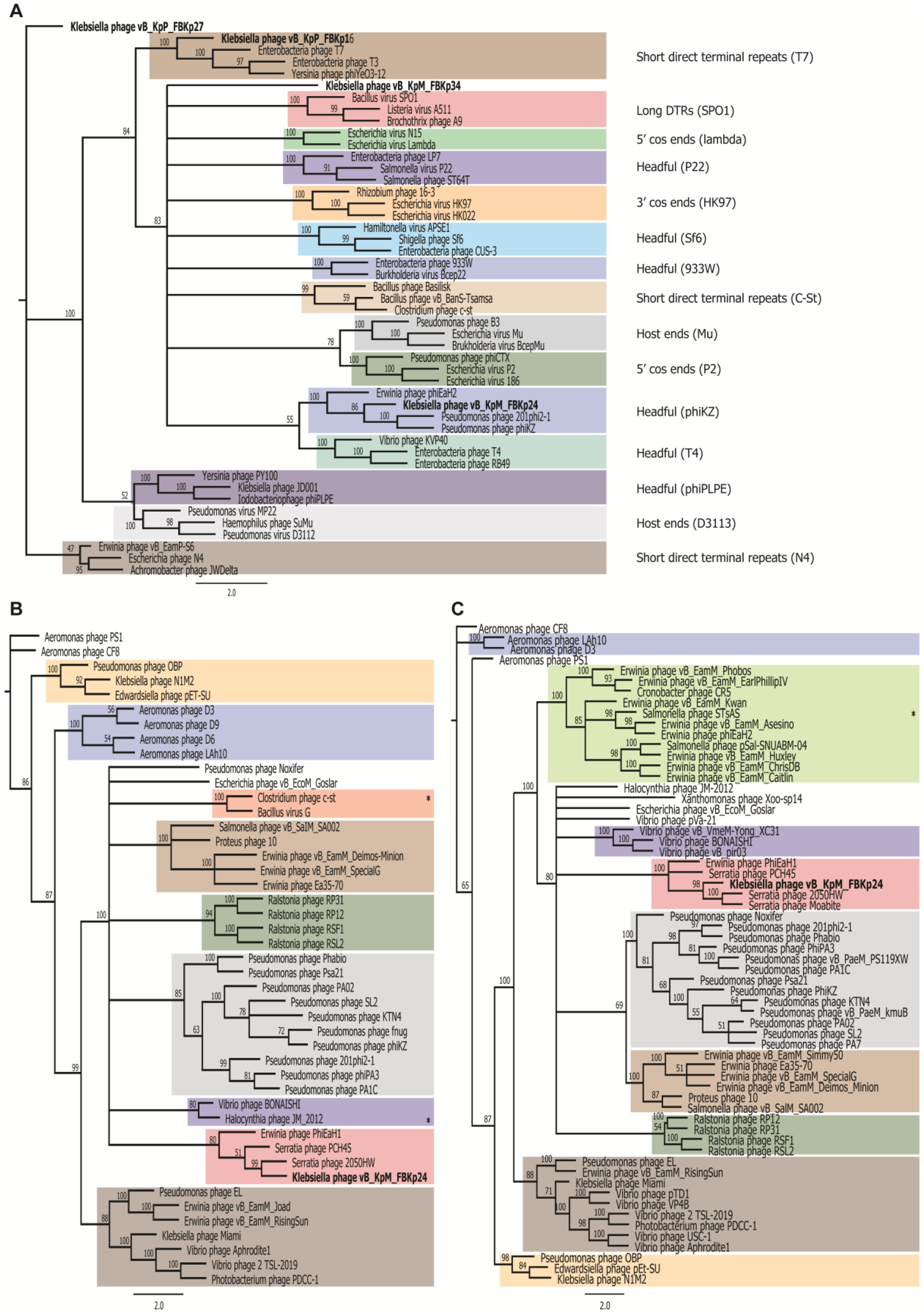
Phylogenetic trees of selected phage proteins. (A) Analysis of large terminase subunits using proteins from phages of well-known packaging mechanisms. (B) Analysis of the tubulin spindle protein of phage ϕKp24 and all protein homologues to the tubulin spindle of phage 201phi2-1 found by psi-Blast and Hmmer. (C) Analysis of the nucleus shell protein of phage ϕKp24 and all protein homologues to the nucleus shell protein of phage 201phi2-1 found by psi-Blast and Hmmer. Trees were built from MAFFT alignments using RAxML with bootstrapping of 100. Identical colors were used in panels (B) and (C) to identify similar phage clusters. All phages in panels (B) and (C) have genomes above 200 kb (Jumbo phages) with the exception of those marked with *, which have a genome size of 167-197 kb.

### 3.4. Bacteriophage ϕKp34 encodes genes with possible anti-viral functions

Phage ϕKp34 encodes an insertion sequence of the IS200/IS605 family (gp188) that is commonly found in bacterial and prophage genomes,^22,52^ suggesting that ϕKp34 can adopt a lysogenic lifestyle as well. IS sequences contribute majorly to bacterial genome diversification, and have also been suggested to play a role in the inactivation and immobilization of other invading phages.^53^ Interestingly, phage ϕKp34 also contains a cluster of genes similar to *terC* (gp124), *terF* C-terminal vWA domain (gp131), and *terD* (gp132, gp133) from the *terZABCDEF* system, and one gene similar to *telA* (gp135) from the *telAB* system. The *terZABCDEF* and *telAB* operons seem to constitute a membrane-linked chemical stress response and anti-viral defense system in bacteria.^54–56^ The subset of genes from the original operons present in ϕKp34 seems to constitute a functional hub found in most major bacterial lineages^56^, has also been reported in virulent phages,^57^ and may confer valuable traits to the bacteria harboring ϕKp34 as a prophage.

### 3.5. Bacteriophage ϕKp24 has multiple depolymerases and tubulin and nuclear shell proteins

Phage ϕKp24 has a distinctive complex structure at its baseplate (**Fig. 1D**) possibly composed of 10 host binding proteins (**Table 1**), in comparison with one host binding protein in phages ϕKp16 and ϕKp34, and three in phage ϕKp27. Three of the 10 possible host binding proteins of ϕKp24 have putative depolymerase domains (**Table 1**) of GTPase, peptidase and transglycosylase activity, while only one host binding protein of ϕKp27 has a predicted depolymerase domain. Depolymerases are used by phages to degrade the capsule of bacteria and to gain access to their secondary receptor (e.g. outer membrane protein, lipopolysaccharide) on the host’s surface. Depolymerases tend to be specific to a capsular type, and the presence of depolymerases with different activities suggests that phage ϕKp24 can interact and degrade different capsular types^46^, likely expanding the phage’s host range.

Interestingly, and akin to other jumbo phages, phage ϕKp24 codes for tubulin spindle (gp094) and nucleus shell (gp083) proteins that function to enhance phage reproduction.^58,59^ The nucleus shell protein forms a proteinaceous barrier that encloses viral DNA, separating phage DNA replication and transcription from other cellular functions and providing a protective physical barrier against DNA-targeting CRISPR-Cas systems;^60,61^ while the tubulin spindle positions the phage nucleus structure at the cell center.^62^ A phylogenetic analysis of the tubulin spindle and nucleus shell proteins of ϕKp24 and all protein homologues to those of phage 201phi2-1 (where these proteins were first reported) found by psi-Blast and Hmmer search (**Fig. 4B** and **4C**) reveals that the proteins of ϕKp24 cluster with those of Serratia phage 2050HW, Serratia phage PCH45, and Erwinia phage PhiEaH1. Interestingly, clusters formed by tubulin spindle and nucleus shell proteins are identical, suggesting that these proteins have co-evolved, and seem to group according to the bacterial species infected. It is also curious that only three of the phages encoding tubulin spindle and nucleus shell proteins have genomes smaller than 200 kb (167-197 kb), further underpinning the exclusive use of these proteins by Jumbo phages.

## 4. Conclusion

The genome sequences of the four *K. pneumoniae* phages reported here reveal potential novel packaging mechanisms (ϕKp27 and ϕKp34), the presence of possible anti-viral strategies in phage genomes (ϕKp34) that can help elucidate symbiotic relationships between temperate phages and their hosts, and distinctive and novel clades of tubulin spindle and nucleus shell proteins (ϕKp24) that will help shed light into the evolution of compartmentalization in prokaryotes and eukaryotes. Phages ϕKp16, ϕKp27 and ϕKp24, but not the potentially temperate phage ϕKp34, are also strong candidates for phage therapy against antibiotic resistant *K. pneumoniae* infections. Further exploration of phage genomes will help elucidate the origins, genetic diversity and evolutionary mechanisms of phages, and contribute to a better understanding of the broader biology of microbial populations and how their genomic characteristics contribute to observable features. This knowledge and the study of individual genes and proteins will certainly also be translated into innovative tools with biotechnological and medical applications.

## Supplementary data

Supplementary data are available at DNARES online.

## Acknowledgements

This work was supported by donations from University Fund from the Delft University of Technology to Fagenbank, as well as generous donations from the public. ARC is supported by the Netherlands Organisation for Scientific Research (NWO) NWA Startimpuls grant 17.366. SJJB is supported by NWO Vici grant VI.C182.027. Genome sequencing and assembly is supported by China National GeneBank and the Global Phage Hub initiated by BGI-Shenzhen. The authors would like to thank Daan Frits van den Berg for the useful discussions about phage tRNAs.

## Author contributions

SJJB, FN, ARC and TvR conceived and designed the project. MX and WS sequenced and assembled the genomes. BEB, ARC, SH and HW annotated the genomes. PJH provided the strains. BEB and ARC generated the data, performed the analysis and wrote the manuscript. SJJB, FN, TvR reviewed and edited the manuscript with input from all authors. All authors approved the manuscript.

## Accession numbers

The assembled and annotated phage genome sequences have been deposited in Genbank (https://www.ncbi.nlm.nih.gov/genbank/) under accession numbers MW394389 (ϕKp16), MW394388 (ϕKp27), MW394391 (ϕKp24) and MW394390 (ϕKp34). The data are also available at the China National GeneBank (CNGB, https://db.cngb.org/) under accession number CNP0000861.

## References

1. Weinbauer, M. G. Ecology of prokaryotic viruses. FEMS Microbiol. Rev. 28, 127–181 (2004).

2. Danovaro, R. et al. Marine viruses and global climate change. FEMS Microbiol. Rev. 35, 993–1034 (2011).

3. Proctor, L. M. & Fuhrman, J. A. Viral mortality of marine bacteria and cyanobacteria. Nature 343, 60–62 (1990).

4. Brüssow, H., Canchaya, C. & Hardt, W.-D. Phages and the Evolution of Bacterial Pathogens: from Genomic Rearrangements to Lysogenic Conversion. Microbiol. Mol. Biol. Rev. 68, 560–602 (2004).

5. Sausset, R., Petit, M. A., Gaboriau-Routhiau, V. & De Paepe, M. New insights into intestinal phages. Mucosal Immunol. 1–11 (2020).

6. Hershey, A. D. & Chase, M. Independent functions of viral protein and nucleic acid in growth of bacteriophage. J. Gen. Physiol. 36, 39–56 (1952).

7. Crick, F. H. C., Barnett, L., Brenner, S. & Watts-Tobin, R. J. General nature of the genetic code for proteins. Nature 192, 1227–1232 (1961).

8. Brenner, S., Jacob, F. & Meselson, M. An unstable intermediate carrying information from genes to ribosomes for protein synthesis. Nature 190, 576–581 (1961).

9. Arber, W. & Linn, S. DNA Modification and Restriction. Annu. Rev. Biochem. 38, 467–500 (1969).

10. Lobban, P. E. & Kaiser, A. D. Enzymatic end-to-end joining of DNA molecules. J. Mol. Biol. 78, 453–471 (1973).

11. Pickar-Oliver, A. & Gersbach, C. A. The next generation of CRISPR–Cas technologies and applications. Nat. Rev. Mol. Cell Biol. 20, 490–507 (2019).

12. O’Neill, J. Tackling drug-resistant infections globally: Final report and recommendations. 2016. HM Gov. Welcome Trust UK (2016).

13. Schroven, K., Aertsen, A. & Lavigne, R. Bacteriophages as drivers of bacterial virulence and their potential for biotechnological exploitation. FEMS Microbiol. Rev. 45, (2021).

14. Pires, D. P., Costa, A. R., Pinto, G., Meneses, L. & Azeredo, J. Current challenges and future opportunities of phage therapy. FEMS Microbiol. Rev. 44, 684–700 (2020).

15. Zampara, A. et al. Developing Innolysins Against Campylobacter jejuni Using a Novel Prophage Receptor-Binding Protein. Front. Microbiol. 12, (2021).

16. Dedrick, R. M. et al. Engineered bacteriophages for treatment of a patient with a disseminated drug-resistant Mycobacterium abscessus. Nat. Med. 25, 730–733 (2019).

17. Tacconelli, E. et al. Discovery, research, and development of new antibiotics: the WHO priority list of antibiotic-resistant bacteria and tuberculosis. Lancet Infect. Dis. 18, 318–327 (2018).

18. Sambrook, J. & Russell, D. W. (David W. Molecular cloning?: a laboratory manual. (Cold Spring Harbor Laboratory Press, 2001).

19. Chen, S., Zhou, Y., Chen, Y. & Gu, J. Fastp: An ultra-fast all-in-one FASTQ preprocessor. In Bioinformatics 34, i884–i890 (Oxford University Press, 2018).

20. Chen, Y. et al. SOAPnuke: A MapReduce acceleration-supported software for integrated quality control and preprocessing of high-throughput sequencing data. Gigascience 7, 1–6 (2018).

21. Li, H. seqtk Toolkit for processing sequences in FASTA/Q formats. GitHub 767, 69 (2012).

22. Bankevich, A. et al. SPAdes: A new genome assembly algorithm and its applications to single-cell sequencing. J. Comput. Biol. 19, 455–477 (2012).

23. Aziz, R. K. et al. The RAST Server: Rapid Annotations using Subsystems Technology. BMC Genomics 9, 75 (2008).

24. Altschul, S. F., Gish, W., Miller, W., Myers, E. W. & Lipman, D. J. Basic local alignment search tool. J. Mol. Biol. 215, 403–410 (1990).

25. Mistry, J., Finn, R. D., Eddy, S. R., Bateman, A. & Punta, M. Challenges in homology search: HMMER3 and convergent evolution of coiled-coil regions. Nucleic Acids Res. 41, (2013).

26. Chan, P. P. & Lowe, T. M. tRNAscan-SE: Searching for tRNA genes in genomic sequences. In Methods in Molecular Biology 1962, 1–14 (Humana Press Inc., 2019).

27. Naville, M., Ghuillot-Gaudeffroy, A., Marchais, A. & Gautheret, D. ARNold: A web tool for the prediction of Rho-independent transcription terminators. RNA Biol. 8, 11–13 (2011).

28. Sampaio, M., Rocha, M., Oliveira, H., Dias, O. & Valencia, A. Predicting promoters in phage genomes using PhagePromoter. Bioinformatics 35, 5301–5302 (2019).

29. Petkau, A., Stuart-Edwards, M., Stothard, P. & van Domselaar, G. Interactive microbial genome visualization with GView. Bioinformatics 26, 3125–3126 (2010).

30. Merrill, B. D., Ward, A. T., Grose, J. H. & Hope, S. Software-based analysis of bacteriophage genomes, physical ends, and packaging strategies. BMC Genomics 17, 1–16 (2016).

31. Rice, P., Longden, I. & Bleasby, A. EMBOSS: the European molecular biology open software suite. Trends Genet. 16, 276–277 (2000).

32. Adriaenssens, E. M. et al. Taxonomy of prokaryotic viruses: 2018-2019 update from the ICTV Bacterial and Archaeal Viruses Subcommittee. Arch. Virol. 165, 1253–1260 (2020).

33. Adriaenssens, E. M. et al. Taxonomy of prokaryotic viruses: 2017 update from the ICTV Bacterial and Archaeal Viruses Subcommittee. Arch. Virol. 163, 1125–1129 (2018).

34. Samson, J. E. & Moineau, S. Characterization of lactococcus lactis phage 949 and comparison with other lactococcal phages. Appl. Environ. Microbiol. 76, 6843–6852 (2010).

35. Dreher, T. W. et al. A freshwater cyanophage whose genome indicates close relationships to photosynthetic marine cyanomyophages. Environ. Microbiol. 13, 1858–1874 (2011).

36. Gervasi, T., Curto, R. Lo, Narbad, A. & Mayer, M. J. Complete genome sequence of ΦCP51, a temperate bacteriophage of Clostridium perfringens. Arch. Virol. 158, 2015–2017 (2013).

37. Bailly-Bechet, M., Vergassola, M. & Rocha, E. Causes for the intriguing presence of tRNAs in phages. Genome Res. 17, 1486–1495 (2007).

38. Kunisawa, T. Functional role of mycobacteriophage transfer RNAs [3]. J. Theor. Biol. 205, 167–170 (2000).

39. Delesalle, V. A., Tanke, N. T., Vill, A. C. & Krukonis, G. P. Testing hypotheses for the presence of tRNA genes in mycobacteriophage genomes. Bacteriophage 6, e1219441 (2016).

40. Eggertsson, G. & Soll, D. Transfer ribonucleic acid-mediated suppression of termination codons in Escherichia coli. Microbiol. Rev. 52, 354–374 (1988).

41. Herring, C. D. & Blattner, F. R. Global transcriptional effects of a suppressor tRNA and the inactivation of the regulator frmR. J. Bacteriol. 186, 6714–6720 (2004).

42. McClain, W. H. UAG suppressor coded by bacteriophage T4. FEBS Lett. 6, 99–101 (1970).

43. Edwards, R. A., McNair, K., Faust, K., Raes, J. & Dutilh, B. E. Computational approaches to predict bacteriophage-host relationships. FEMS Microbiol. Rev. 40, 258–272 (2016).

44. Marinelli, L. J. et al. Propionibacterium acnes bacteriophages display limited genetic diversity and broad killing activity against bacterial skin isolates. MBio 3, (2012).

45. Simoliunas, E. et al. Genome of Klebsiella sp.-Infecting Bacteriophage vB_KleM_RaK2. J. Virol. 86, 5406–5406 (2012).

46. Pan, Y.-J. et al. Klebsiella Phage ΦK64-1 Encodes Multiple Depolymerases for Multiple Host Capsular Types. J. Virol. 91, e02457–16 (2017).

47. Dupuis M. È. & Moineau, S. Genome organization and characterization of the virulent lactococcal phage 1358 and its similarities to Listeria phages. Appl. Environ. Microbiol. 76, 1623–1632 (2010).

48. Leptihn, S., Gottschalk, J. & Kuhn, A. T7 ejectosome assembly: A story unfolds. Bacteriophage 6, e1128513 (2016).

49. Wangchuk, J., Prakash, P., Bhaumik, P. & Kondabagil, K. Bacteriophage N4 large terminase: Expression, purification and X-ray crystallographic analysis. Acta Crystallogr. Sect. F Struct. Biol. Commun. 74, 198–204 (2018).

50. Buttimer, C. et al. Novel N4-like bacteriophages of pectobacterium atrosepticum. Pharmaceuticals 11, (2018).

51. Shi, X. et al. Characterization and Complete Genome Analysis of Pseudomonas aeruginosa Bacteriophage vB_PaeP_LP14 Belonging to Genus Litunavirus. Curr. Microbiol. 77, 2465–2474 (2020).

52. Kuno, S., Yoshida, T., Kamikawa, R., Hosoda, N. & Sako, Y. The distribution of a phage-related insertion sequence element in the cyanobacterium, Microcystis aeruginosa. Microbes Environ. 25, 295–301 (2010).

53. Ooka, T. et al. Inference of the impact of insertion sequence (IS) elements on bacterial genome diversification through analysis of small-size structural polymorphisms in Escherichia coli O157 genomes. Genome Res. 19, 1809–1816 (2009).

54. Whelan, K. F., Colleran, E. & Taylor, D. E. Phage inhibition, colicin resistance, and tellurite resistance are encoded by a single cluster of genes on the IncHI2 plasmid R478. J. Bacteriol. 177, 5016–5027 (1995).

55. Walter, E. G., Thomas, C. M., Ibbotson, J. P. & Taylor, D. E. Transcriptional analysis, translational analysis, and sequence of the kilA-tellurite resistance region of plasmid RK2Te(r). J. Bacteriol. 173, 1111–1119 (1991).

56. Anantharaman, V., Iyer, L. M. & Aravind, L. Ter-dependent stress response systems: Novel pathways related to metal sensing, production of a nucleoside-like metabolite, and DNA-processing. Mol. Biosyst. 8, 3142–3165 (2012).

57. Frampton, R. A. et al. Identification of bacteriophages for biocontrol of the kiwifruit canker phytopathogen Pseudomonas syringae pv. actinidiae. Appl. Environ. Microbiol. 80, 2216–2228 (2014).

58. Kraemer, J. A. et al. A phage tubulin assembles dynamic filaments by an atypical mechanism to center viral DNA within the host cell. Cell 149, 1488–1499 (2012).

59. Erb, M. L. et al. A bacteriophage tubulin harnesses dynamic instability to center DNA in infected cells. Elife 3, (2014).

60. Mendoza, S. D. et al. A bacteriophage nucleus-like compartment shields DNA from CRISPR nucleases. Nature 577, 244–248 (2020).

61. Malone, L. M. et al. A jumbo phage that forms a nucleus-like structure evades CRISPR–Cas DNA targeting but is vulnerable to type III RNA-based immunity. Nat. Microbiol. 5, 48–55 (2020).

62. Chaikeeratisak, V. et al. The Phage Nucleus and Tubulin Spindle Are Conserved among Large Pseudomonas Phages. Cell Rep. 20, 1563–1571 (2017).

