## Supplementary Table for "Genomic characterization of four novel bacteriophages infecting the clinical pathogen *Klebsiella pneumoniae*"

### Content

**Supplementary Table S1.** List of protein homologs to the tubulin spindle protein of phage 201phi2-1 found by psi-Blast and Hmmer search and used to build the tubulin spindle phylogenetic tree.

**Supplementary Table S2.** List of protein homologs to the nucleus shell protein of phage 201phi2-1 found by psi-Blast and Hmmer search and used to build the nucleus shell phylogenetic tree.

**Supplementary Table S3.** Genome annotations of *Klebsiella pneumoniae* phage vB_KpP_FBKp16.

**Supplementary Table S4.** Genome annotations of *Klebsiella pneumoniae* phage vB_KpP_FBKp27.

**Supplementary Table S5.** Genome annotations of *Klebsiella pneumoniae* phage vB_KpM_FBKp34.

**Supplementary Table S6.** Genome annotations of *Klebsiella pneumoniae* phage vB_KpM_FBKp24.

**Supplementary Table S7.** Analysis of codon usage of phages vB_KpP_FBKp27, vB_KpM_FBKp34, vB_KpM_FBKp24 and *Klebsiella pneumoniae* HS11286 using Cusp from EMBOSS.

**Supplementary Table S1.** List of protein homologs to the tubulin spindle protein of phage 201phi2-1 found by psi-Blast and Hmmer search and used to build the tubulin spindle phylogenetic tree.

| **Phage** | **Genome size** | **psi-Blast** | | | **Hmmer E value** | **Protein accession number** | **Ref** |
|---|---|---|---|---|---|---|---|
| | | **Coverage (%)** | **E-value** | **Identity (%)** | | | |
| *Aeromonas* phage CF8 | 238,150 | 100 | 5.0E-55 | 24.46 | - | QDB70454.1 | - |
| *Aeromonas* phage D3 | 262,372 | 98 | 6.0E-50 | 18.61 | - | QDJ97033.1 | - |
| *Aeromonas* phage D6 | 259,831 | 98 | 5.0E-49 | 18.93 | - | QDJ97195.1 | - |
| *Aeromonas* phage D9 | 260,857 | 98 | 6.0E-50 | 18.61 | - | QEP52339.1 | - |
| *Aeromonas* phage LAh10 | 260,310 | 98 | 5.0E-49 | 18.93 | - | QDH47086.1 | - |
| *Aeromonas* phage PS1 | 237,367 | 99 | 1.0E-47 | 20.56 | - | QDJ96797.1 | - |
| *Bacillus* virus G | 497,513 | 94 | 7.0E-53 | 12.28 | - | YP_009015441.1 | - |
| *Clostridium* phage c-st | 185,683 | 95 | 4.0E-65 | 15.00 | - | 4XCQ_A | - |
| *Edwardsiella* phage pEt-SU | 276,734 | 99 | 2.0E-58 | 25.39 | 3.0E-18 | YP_009822177.1 | - |
| *Erwinia* phage Ea35-70 | 271,084 | 100 | 3.0E-56 | 23.46 | 1.0E-21 | YP_009005002.1 | - |
| *Erwinia* phage PhiEaH1 | 218,339 | 98 | 7.0E-49 | 23.84 | 2.6E-15 | YP_009010087.1 | - |
| *Erwinia* phage vB_EamM_Deimos-Minion | 273,501 | - | - | - | 1.0E-21 | ANH52315 | - |
| *Erwinia* phage vB_EamM_Joad | 235,374 | - | - | - | 1.2E-06 | ASU03898 | - |
| *Erwinia* phage vB_EamM_RisingSun | 235,108 | 99 | 2.0E-48 | 16.92 | - | YP_009612761.1 | - |
| *Erwinia* phage vB_EamM_Special G | 273,224 | - | - | - | 1.0E-21 | ANJ65024 | - |
| *Escherichia* phage vB_EcoM_Goslar | 237,307 | 99 | 2.0E-55 | 22.77 | 1.3E-12 | YP_009820885.1 | - |
| *Halocynthia* phage JM-2012 | 167,292 | 99 | 2.0E-43 | 20.81 | 6.3E-09 | YP_006383382.1 | - |
| *Klebsiella* phage Miami | 253,383 | 98 | 2.0E-44 | 18.81 | - | QPB09366.1 | - |
| *Klebsiella* phage N1M2 | 253,367 | 99 | 8.0E-60 | 21.81 | - | QGH71890.1 | - |
| *Photobacterium* phage PDCC-1 | 237,509 | 98 | 6.0E-52 | 19.44 | - | YP_009853560.1 | - |
| *Proteus* phage 10 | 223,209 | 100 | 6.0E-61 | 25.00 | - | QMP24134.1 | - |
| *Pseudomonas* phage 201phi2-1 | 316,674 | 99 | 8.0E-57 | 100.00 | 5.1E-208 | 3J5V_a | [1] |
| *Pseudomonas* phage EL | 211,215 | 91 | 3.0E-45 | 16.72 | - | YP_418049.1 | - |
| *Pseudomonas* phage fnug | 278,899 | 99 | 3.0E-63 | 29.85 | - | QJB22689.1 | - |
| *Pseudomonas* phage KTN4 | 279,593 | 99 | 2.0E-63 | 29.85 | - | ANM44810.1 | - |
| *Pseudomonas* phage Noxifer | 278,136 | 99 | 4.0E-46 | 26.93 | 1.9E-26 | YP_009608946.1 | - |
| *Pseudomonas* phage OBP | 284,757 | 99 | 3.0E-49 | 22.12 | 2.0E-14 | YP_004957954.1 | - |
| *Pseudomonas* phage PA02 | 279,095 | 99 | 2.0E-64 | 30.46 | - | BBI55877.1 | - |
| *Pseudomonas* phage PA1C | 304,671 | 99 | 2.0E-76 | 48.73 | 7.1E-88 | QBX32179.1 | - |
| *Pseudomonas* phage Phabio | 309,157 | 99 | 6.0E-52 | 29.84 | 2.7E-39 | ARV76677.1 | - |
| *Pseudomonas* phage PhiPA3 | 309,208 | 99 | 6.0E-73 | 45.86 | 2.0E-86 | YP_009217111.1 | [1] |

| | | | | | | | |
|---|---|---|---|---|---|---|---|
| *Pseudomonas* phage Psa21 | 305,260 | 99 | 8.0E-54 | 31.95 | 1.5E-46 | QBJ02557.1 | - |
| *Pseudomonas* phage SL2 | 279,696 | 99 | 3.0E-63 | 29.85 | - | YP_009619864.1 | - |
| *Pseudomonas* virus phiKZ | 278,899 | 99 | 2.0E-63 | 29.85 | 2.8E-44 | 3ZBP_A | [1] |
| *Ralstonia* phage RP12 | 279,845 | 96 | 3.0E-51 | 19.62 | 2.0E-09 | YP_009598707.1 | - |
| *Ralstonia* phage RP31 | 276,958 | 96 | 2.0E-51 | 19.94 | 1.9E-09 | BAW19276.1 | - |
| *Ralstonia* phage RSF1 | 222,888 | 91 | 5.0E-48 | 20.76 | 1.3E-12 | YP_009207815.1 | - |
| *Ralstonia* phage RSL2 | 223,932 | 91 | 9.0E-51 | 22.15 | 1.1E-13 | BAQ02539.2 | - |
| *Salmonella* phage vB_SalM_SA002 | 288,012 | 100 | 9.0E-59 | 24.54 | - | QKE54534.1 | - |
| *Serratia* phage 2050HW | 276,025 | 99 | 2.0E-56 | 22.50 | 1.6E-16 | YP_009833975.1 | [2] |
| *Serratia* phage PCH45 | 212,807 | 90 | 2.0E-51 | 18.47 | - | QFP93107.1 | - |
| *Vibrio* phage 2 TSL-2019 | 242,446 | 98 | 2.0E-51 | 19.44 | - | YP_009843109.1 | - |
| *Vibrio* phage Aphrodite1 | 237,722 | 98 | 8.0E-54 | 21.25 | 3.5E-13 | YP_009622183.1 | - |
| *Vibrio* phage BONAISHI | 288,967 | 98 | 2.0E-49 | 20.24 | 1.7E-11 | AXH70800.1 | - |

**Supplementary Table S2.** List of protein homologs to the nucleus shell protein of phage 201phi2-1 found by psi-Blast and Hmmer search and used to build the nucleus shell phylogenetic tree.

| Phage | Genome size | psi-Blast | | | Hmmer | Protein accession number | Ref |
|---|---|---|---|---|---|---|---|
| | | Coverage (%) | E-value | Identity (%) | E value | | |
| *Aeromonas* phage CF8 | 238,150 | 97 | 5.0E-77 | 14.01 | - | QDB70465.1 | - |
| *Aeromonas* phage D3 | 262,372 | 96 | 6.0E-90 | 13.73 | - | QDJ97041.1 | - |
| *Aeromonas* phage LAh10 | 260,310 | 96 | 1.0E-89 | 13.27 | - | QDH47031.1 | - |
| *Aeromonas* phage PS1 | 237,367 | 93 | 3.0E-94 | 13.90 | - | QDJ96807.1 | - |
| *Cronobacter* phage CR5 | 223,989 | 95 | 2.0E-96 | 13.76 | - | YP_008125763.1 | - |
| *Edwardsiella* phage pEt-SU | 276,734 | 92 | 2.0E-96 | 13.29 | - | YP_009821902.1 | - |
| *Erwinia* phage Ea35-70 | 271,084 | 96 | 1.0E-116 | 20.79 | 3.6E-12 | YP_009005014.1 | - |
| *Erwinia* phage PhiEaH1 | 218,339 | 98 | 7.0E-91 | 16.25 | - | YP_009010075.1 | - |
| *Erwinia* phage phiEaH2 | 243,050 | 90 | 5.0E-115 | 12.48 | - | YP_007237854.1 | - |
| *Erwinia* phage vB_EamM_Asesino | 246,290 | 99 | 7.0E-117 | 11.85 | - | YP_009290657.1 | - |
| *Erwinia* phage vB_EamM_Caitlin | 241,147 | 90 | 3.0E-104 | 12.88 | - | YP_009292105.1 | - |
| *Erwinia* phage vB_EamM_ChrisDB | 244,840 | 89 | 4.0E-103 | 13.14 | - | YP_009292727.1 | - |
| *Erwinia* phage vB_EamM_Deimos-Minion | 273,501 | - | - | - | 3.6E-12 | ANH52327 | - |
| *Erwinia* phage vB_EamM_EarlPhillipIV | 223,935 | 95 | 8.0E-96 | 13.85 | - | YP_009278336.1 | - |
| *Erwinia* phage vB_EamM_Huxley | 240,761 | 96 | 5.0E-105 | 11.73 | - | YP_009293007.1 | - |
| *Erwinia* phage vB_EamM_Kwan | 246,390 | 90 | 2.0E-91 | 12.01 | - | YP_009278639.1 | - |
| *Erwinia* phage vB_EamM_Phobos | 229,501 | 95 | 3.0E-94 | 13.76 | - | YP_009283513.1 | - |
| *Erwinia* phage vB_EamM_RisingSun | 235,108 | 94 | 7.0E-76 | 12.98 | - | YP_009612772.1 | - |
| *Erwinia* phage vB_EamM_Simmy50 | 271,088 | 93 | 5.0E-116 | 21.39 | - | YP_009606010.1 | - |
| *Erwinia* phage vB_EamM_Special G | 273,224 | - | - | - | 3.6E-12 | ANJ65036 | - |
| *Escherichia* phage vB_EcoM_Goslar | 237,307 | 100 | 2.0E-131 | 18.18 | 2.8E-07 | YP_009820873.1 | - |
| *Halocynthia* phage JM-2012 | 167,292 | 94 | 4.0E-55 | 14.77 | - | YP_006383413.1 | - |
| *Klebsiella* phage Miami | 253,383 | 95 | 5.0E-80 | 15.76 | - | QPB09376.1 | - |
| *Klebsiella* phage N1M2 | 253,367 | 92 | 7.0E-95 | 15.24 | - | QGH71899.1 | - |
| *Photobacterium* phage PDCC-1 | 237,509 | 99 | 5.0E-91 | 15.06 | - | YP_009853356.1 | - |

| | | | | | | | |
|---|---|---|---|---|---|---|---|
| *Proteus* phage 10 | 223,209 | 97 | 3.0E-113 | 20.60 | - | QMP24145.1 | - |
| *Pseudomonas* phage 201phi2-1 | 316,674 | 100 | 2.0E-175 | 100.00 | 0.0E+00 | YP_001956829.1 | [1] |
| *Pseudomonas* phage EL | 211,215 | 97 | 2.0E-68 | 11.11 | - | YP_418056.1 | - |
| *Pseudomonas* phage KTN4 | 279,593 | 100 | 2.0E-146 | 39.59 | - | ANM44831.1 | - |
| *Pseudomonas* phage Noxifer | 278,136 | 96 | 3.0E-142 | 35.90 | 1.7E-111 | YP_009608965.1 | - |
| *Pseudomonas* phage OBP | 284,757 | 93 | 4.0E-102 | 14.38 | - | YP_004957964.1 | - |
| *Pseudomonas* phage PA02 | 279,095 | 100 | 2.0E-145 | 39.59 | - | BBI55859.1 | - |
| *Pseudomonas* phage PA1C | 304,671 | 100 | 1.0E-164 | 51.35 | 1.1E-201 | QBX32206.1 | - |
| *Pseudomonas* phage PA7 | 266,743 | 100 | 2.0E-145 | 39.44 | - | YP_009617618.1 | - |
| *Pseudomonas* phage Phabio | 309,157 | 100 | 1.0E-156 | 52.52 | 7.8E-206 | ARV76713.1 | - |
| *Pseudomonas* phage PhiPA3 | 309,208 | 95 | 5.0E-172 | 51.41 | 6.2E-196 | YP_009217136.1 | [1] |
| *Pseudomonas* phage Psa21 | 305,260 | 100 | 3.0E-139 | 41.13 | 4.8E-144 | QBJ02586.1 | - |
| *Pseudomonas* phage SL2 | 279,696 | 100 | 3.0E-145 | 39.28 | - | YP_009619844.1 | - |
| *Pseudomonas* phage vB_PaeM_kmuB | 266,592 | 100 | 1.0E-146 | 39.59 | - | QOV07924.1 | - |
| *Pseudomonas* phage vB_PaeM_PS119XW | 301,543 | 100 | 2.0E-166 | 51.35 | - | QEM41783.1 | - |
| *Pseudomonas* virus phiKZ | 278,899 | 100 | 5.0E-146 | 39.75 | 2.1E-133 | NP_803620.1 | [1] |
| *Ralstonia* phage RP12 | 279,845 | 99 | 1.0E-95 | 18.88 | 8.1E-15 | YP_009598981.1 | - |
| *Ralstonia* phage RP31 | 276,958 | 99 | 5.0E-93 | 18.82 | 5.2E-15 | BAW19549.1 | - |
| *Ralstonia* phage RSF1 | 222,888 | 97 | 2.0E-118 | 17.67 | 6.9E-07 | YP_009208033.2 | - |
| *Ralstonia* phage RSL2 | 223,932 | 97 | 9.0E-117 | 17.80 | 1.5E-07 | YP_009213072.1 | - |
| *Salmonella* phage pSal-SNUABM-04 | 239,626 | 94 | 2.0E-103 | 11.49 | - | QOC54489.1 | - |
| *Salmonella* phage STsAS | 197,215 | 99 | 2.0E-108 | 13.64 | - | AWN08967.1 | - |
| *Salmonella* phage vB_SalM_SA002 | 288,012 | 95 | 6.0E-120 | 20.29 | - | QKE54546.1 | - |
| *Serratia* phage 2050HW | 276,025 | 37 | 1.0E-21 | 14.57 | - | YP_009833990.1 | [2] |
| *Serratia* phage Moabite | 273,933 | 90 | 6.0E-84 | 16.47 | - | YP_009849136.1 | - |
| *Serratia* phage PCH45 | 212,807 | 95 | 6.0E-85 | 13.94 | - | QFP93061.1 | - |
| *Vibrio* phage 2 TSL-2019 | 242,446 | 99 | 7.0E-90 | 15.06 | - | YP_009843316.1 | - |
| *Vibrio* phage Aphrodite1 | 237,722 | 99 | 2.0E-87 | 15.21 | - | YP_009622192.1 | - |
| *Vibrio* phage BONAISHI | 288,967 | 99 | 3.0E-88 | 14.85 | - | AXH70812.1 | - |
| *Vibrio* phage pTD1 | 239,276 | 94 | 8.0E-92 | 14.93 | - | YP_009599417.1 | - |
| *Vibrio* phage pVa-21 | 231,998 | 93 | 4.0E-82 | 13.79 | - | AQT28149.1 | - |
| *Vibrio* phage USC-1 | 238,099 | 99 | 2.0E-87 | 15.21 | - | YP_009847864.1 | - |
| *Vibrio* phage vB_pir03 | 286,284 | 98 | 2.0E-70 | 15.85 | - | QNI21050.1 | - |
| *Vibrio* phage vB_VmeM-Yong XC31 | 290,532 | 97 | 2.0E-78 | 15.24 | - | QAX96061.1 | - |
| *Vibrio* phage VP4B | 236,053 | 99 | 5.0E-96 | 16.01 | - | YP_009626053.1 | - |
| *Xanthomonas* phage Xoo-sp14 | 232,104 | 99 | 1.0E-105 | 19.82 | - | QNR51901.1 | - |

**Supplementary Table S3.** Genome annotations of *Klebsiella pneumoniae* phage vB_KpP_FBKp16.

| Locus_tag | Direction | Min | Max | Product | Notes | Function |
|---|---|---|---|---|---|---|
| gp001 | + | 13 | 729 | hypothetical protein | | |
| gp002 | + | 739 | 1,086 | peptidase | contains peptidase M15 domain | cell lysis |
| gp003 | + | 1,079 | 1,201 | hypothetical protein | | |
| gp004 | + | 1,198 | 1,344 | hypothetical protein | | |
| gp005 | + | 1,413 | 1,628 | hypothetical protein | | |
| gp006 | + | 1,949 | 3,694 | hypothetical protein | | |
| gp007 | + | 5,005 | 5,205 | hypothetical protein | | |
| gp008 | - | 5,479 | 5,709 | hypothetical protein | | |
| gp009 | + | 5,774 | 5,968 | hypothetical protein | | |
| gp010 | + | 6,023 | 6,541 | hypothetical protein | | |
| gp011 | + | 6,737 | 7,642 | hypothetical protein | | |
| gp012 | + | 7,711 | 10,344 | phage DNA-directed RNA polymerase | contains DNA-directed RNA polymerase N-terminal and DNA-dependent RNA polymerase domains | transcription |
| gp013 | + | 10,594 | 10,770 | hypothetical protein | | |
| gp014 | + | 10,773 | 12,776 | DNA primase/helicase | DnaB-like helicase C terminal domain | DNA replication |
| gp015 | + | 12,840 | 13,535 | hypothetical protein | | |
| gp016 | + | 13,532 | 13,759 | hypothetical protein | | |
| gp017 | + | 13,752 | 13,964 | hypothetical protein | | |
| gp018 | + | 14,030 | 14,125 | hypothetical protein | | |
| gp019 | + | 14,109 | 16,655 | phage DNA-directed DNA polymerase | contains DNA polymerase family A domain | DNA replication |
| gp020 | + | 16,666 | 16,887 | hypothetical protein | | |
| gp021 | + | 17,026 | 17,832 | hypothetical protein | | |
| gp022 | + | 17,832 | 18,047 | hypothetical protein | | |
| gp023 | + | 18,345 | 18,716 | hypothetical protein | | |

| Locus_tag | Direction | Min | Max | Product | Notes | Function |
|---|---|---|---|---|---|---|
| gp024 | + | 18,796 | 19,242 | hypothetical protein | | |
| gp025 | + | 19,310 | 19,633 | hypothetical protein | | |
| gp026 | + | 19,615 | 20,550 | phage exonuclease | | DNA replication |
| gp027 | + | 20,535 | 20,936 | endonuclease | contains recombination endonuclease VII domain | DNA repair |
| gp028 | + | 20,937 | 21,944 | phage phosphoesterase | | |
| gp029 | + | 22,061 | 22,468 | hypothetical protein | | |
| gp030 | + | 22,465 | 22,668 | hypothetical protein | | |
| gp031 | + | 22,665 | 23,618 | phage-associated ATP-dependent DNA ligase | contains DNA ligase OB-like domain | DNA replication and repair |
| gp032 | + | 23,611 | 23,796 | hypothetical protein | | |
| gp033 | + | 23,783 | 24,211 | HNH endonuclease | | DNA metabolism |
| gp034 | + | 24,238 | 24,387 | hypothetical protein | | |
| gp035 | + | 24,384 | 24,845 | putative N-acetyltransferase | contains acetyltransferase (GNAT) family domain | |
| gp036 | + | 24,854 | 25,060 | hypothetical protein | | |
| gp037 | + | 25,062 | 26,630 | phage collar, head-to-tail connector protein | contains bacteriophage head to tail connecting protein domain | structural components |
| gp038 | + | 26,630 | 27,481 | phage capsid assembly scaffolding protein | | structural components |
| gp039 | + | 27,557 | 28,648 | major capsid protein | | structural components |
| gp040 | + | 28,800 | 29,525 | phage non-contractile tail tubular protein A | contains tail tubular protein domain | structural components |
| gp041 | + | 29,525 | 31,909 | phage non-contractile tail tubular protein B | | structural components |
| gp042 | + | 31,909 | 32,577 | internal virion protein | | structural components |
| gp043 | + | 32,578 | 35,514 | hypothetical protein | | |
| gp044 | + | 35,584 | 39,396 | phage DNA ejectosome component Gp16, peptidoglycan lytic exotransglycosylase (EC 4.2.2.n1) | | structural components |

| Locus_tag | Direction | Min | Max | Product | Notes | Function |
|---|---|---|---|---|---|---|
| gp045 | + | 39,396 | 40,382 | tail fiber | contains phage T7 tail fibre protein domain; coiledstalk of trimeric autotransporter adhesin domain | structural components |
| gp046 | + | 40,391 | 40,579 | holin | contains phage holin T7 family, holin superfamily II domain | cell lysis |
| gp047 | + | 40,569 | 40,868 | phage terminase, small subunit | | DNA packaging |
| gp048 | + | 40,868 | 42,766 | phage terminase large subunit | | |
| gp049 | + | 42,917 | 43,183 | hypothetical protein | | |
| gp050 | + | 43,197 | 43,463 | hypothetical protein | | |
| gp051 | + | 43,474 | 44,010 | hypothetical protein | possible neck whiskers protein | |

**Supplementary Table S4.** Genome annotations of *Klebsiella pneumoniae* phage vB_KpP_FBKp27.

| Locus_tag | Direction | Min | Max | Product | Notes | Function | Cluster |
|---|---|---|---|---|---|---|---|
| gp001 | - | 46 | 744 | hypothetical protein | putative tail protein | structural components | F |
| gp002 | + | 834 | 1,595 | thymidylate synthase thyX | | DNA metabolism | A |
| gp003 | + | 1,683 | 2,747 | ribonucleotide reductase, small subunit | | DNA metabolism | A |
| gp004 | + | 2,750 | 4,480 | ribonucleotide reductase of class Ia (aerobic), alpha subunit | | DNA metabolism | A |
| gp005 | + | 4,538 | 5,017 | hypothetical protein | | | A |
| gp006 | + | 4,995 | 5,969 | hypothetical protein | | | A |
| gp007 | + | 5,978 | 8,137 | hypothetical protein | DNA primase | | A |
| gp008 | + | 8,207 | 8,938 | hypothetical protein | AAA-domain protein | | A |
| gp009 | + | 8,935 | 9,465 | putative HNH endonuclease | | DNA metabolism | A |
| gp010 | + | 9,467 | 10,207 | hypothetical protein | putative single-stranded DNA-binding protein | | A |
| gp011 | + | 10,250 | 10,573 | hypothetical protein | | | A |
| gp012 | + | 10,536 | 11,099 | hypothetical protein | | | A |
| gp013 | + | 11,102 | 11,782 | hypothetical protein | | | A |
| gp014 | + | 11,793 | 11,966 | hypothetical protein | | | A |
| gp015 | - | 12,002 | 22,642 | RNA polymerase | | transcription | B |
| gp016 | - | 22,687 | 23,745 | hypothetical protein | | | B |
| gp017 | - | 23,749 | 24,174 | hypothetical protein | | | B |
| gp018 | - | 24,183 | 26,897 | hypothetical protein | | | B |
| gp019 | - | 26,954 | 27,772 | hypothetical protein | | | B |
| gp020 | - | 27,782 | 28,324 | hypothetical protein | | | B |
| gp021 | - | 28,408 | 29,580 | major capsid protein | | structural components | B |
| gp022 | - | 29,595 | 30,755 | putative tail tape measure protein | | structural components | B |
| gp023 | - | 30,759 | 31,073 | hypothetical protein | | | B |

| Locus_tag | Direction | Min | Max | Product | Notes | Function | Cluster |
|---|---|---|---|---|---|---|---|
| gp024 | - | 31,143 | 33,404 | portal protein | | structural components | B |
| gp025 | + | 33,461 | 34,075 | putative nucleoside triphosphate pyrophosphohydrolase | | regulation | C |
| tRNA | + | 34,455 | 34,529 | tRNA-Asn-GTT | | | C |
| tRNA | + | 34,535 | 34,609 | tRNA-Asp-GTC | | | C |
| tRNA | + | 34,617 | 34,693 | tRNA-Pro-TGG | | | C |
| tRNA | + | 34,699 | 34,785 | tRNA-Tyr-GTA | | | C |
| tRNA | + | 34,792 | 34,881 | tRNA-Ser-TGA | | | C |
| tRNA | + | 34,955 | 35,031 | tRNA-Gln-TTG | | | C |
| gp026 | + | 35,311 | 35,622 | hypothetical protein | | | C |
| gp027 | + | 35,619 | 36,038 | hypothetical protein | | | C |
| gp028 | - | 36,068 | 36,949 | hypothetical protein | | | D |
| gp029 | - | 36,959 | 38,563 | terminase large subunit | | DNA packaging | D |
| gp030 | - | 38,665 | 39,360 | hypothetical protein | | | D |
| gp031 | + | 39,445 | 39,840 | hypothetical protein | | | D |
| gp032 | - | 39,873 | 40,067 | hypothetical protein | | | D |
| gp033 | + | 40,591 | 40,818 | hypothetical protein | | | E |
| gp034 | + | 41,445 | 41,588 | hypothetical protein | | | E |
| gp035 | + | 41,596 | 41,748 | hypothetical protein | | | E |
| gp036 | + | 41,752 | 41,943 | hypothetical protein | | | E |
| gp037 | + | 41,959 | 42,222 | hypothetical protein | | | E |
| gp038 | + | 42,264 | 42,536 | hypothetical protein | | | E |
| gp039 | + | 42,816 | 43,019 | hypothetical protein | | | E |
| gp040 | + | 43,006 | 43,245 | hypothetical protein | | | E |
| gp041 | + | 43,666 | 44,007 | hypothetical protein | | | E |
| gp042 | + | 44,007 | 44,135 | hypothetical protein | | | E |

| Locus_tag | Direction | Min | Max | Product | Notes | Function | Cluster |
|---|---|---|---|---|---|---|---|
| gp043 | + | 44,137 | 44,391 | hypothetical protein | | | E |
| gp044 | + | 44,378 | 44,671 | hypothetical protein | | | E |
| gp045 | + | 44,668 | 44,928 | hypothetical protein | | | E |
| gp046 | + | 45,011 | 45,301 | hypothetical protein | | | E |
| gp047 | + | 45,350 | 45,661 | hypothetical protein | | | E |
| gp048 | + | 45,651 | 45,929 | hypothetical protein | | | E |
| gp049 | + | 45,929 | 46,936 | RNA polymerase small subunit | | transcription | E |
| gp050 | + | 46,954 | 47,187 | hypothetical protein | | | E |
| gp051 | + | 47,177 | 48,451 | RNA polymerase large subunit | | transcription | E |
| gp052 | + | 48,438 | 48,572 | hypothetical protein | | | E |
| gp053 | + | 48,618 | 48,839 | hypothetical protein | | | E |
| gp054 | + | 48,830 | 49,078 | hypothetical protein | | | E |
| gp055 | + | 49,084 | 49,473 | hypothetical protein | | | E |
| gp056 | + | 49,525 | 49,809 | hypothetical protein | | | E |
| gp057 | + | 50,361 | 50,609 | hypothetical protein | | | E |
| gp058 | + | 50,602 | 50,763 | hypothetical protein | | | E |
| gp059 | + | 50,738 | 51,400 | hypothetical protein | | | E |
| gp060 | + | 51,421 | 51,591 | hypothetical protein | | | E |
| gp061 | + | 51,588 | 52,130 | hypothetical protein | | | E |
| gp062 | + | 52,114 | 52,302 | hypothetical protein | | | E |
| gp063 | + | 52,265 | 52,630 | putative HNH endonuclease | | DNA metabolism | E |
| gp064 | + | 52,627 | 52,959 | hypothetical protein | | | E |
| gp065 | + | 52,943 | 53,128 | hypothetical protein | | | E |
| gp066 | + | 53,125 | 53,298 | hypothetical protein | | | E |
| gp067 | + | 53,355 | 53,552 | hypothetical protein | | | E |

| Locus_tag | Direction | Min | Max | Product | Notes | Function | Cluster |
|---|---|---|---|---|---|---|---|
| gp068 | + | 53,555 | 54,226 | hypothetical protein | | | E |
| gp069 | + | 54,236 | 54,475 | hypothetical protein | | | E |
| gp070 | + | 54,477 | 55,529 | putative ATPase | | DNA replication | E |
| gp071 | + | 55,539 | 56,720 | hypothetical protein | | | E |
| gp072 | + | 56,859 | 58,142 | DNA helicase | | DNA replication | E |
| gp073 | + | 58,139 | 58,660 | hypothetical protein | | | E |
| gp074 | + | 58,657 | 59,031 | hypothetical protein | | | E |
| gp075 | + | 59,031 | 59,399 | hypothetical protein | | | E |
| gp076 | + | 59,356 | 59,535 | hypothetical protein | | | E |
| gp077 | + | 59,606 | 62,083 | RIIA-like protein | | DNA replication | E |
| gp078 | + | 62,080 | 63,930 | RIIB-like protein | | DNA replication | E |
| gp079 | + | 63,983 | 66,607 | DNA polymerase I | | DNA replication | E |
| gp080 | + | 66,600 | 67,100 | hypothetical protein | putative avirulence protein | | E |
| gp081 | + | 67,094 | 67,468 | hypothetical protein | | | E |
| gp082 | + | 67,513 | 67,836 | hypothetical protein | | | E |
| gp083 | + | 67,877 | 68,215 | hypothetical protein | | | E |
| gp084 | + | 68,232 | 69,131 | methyl-directed repair DNA adenine methylase | | DNA repair | E |
| gp085 | + | 69,128 | 69,358 | hypothetical protein | | | E |
| gp086 | - | 69,583 | 70,026 | putative tail protein | | structural components | F |
| gp087 | - | 70,103 | 70,612 | endopeptidase Rz | | cell lysis | F |
| gp088 | - | 70,593 | 70,814 | hypothetical protein | possible holin | | F |
| gp089 | - | 70,811 | 71,299 | lysozyme | | cell lysis | F |
| gp090 | - | 71,339 | 71,533 | hypothetical protein | | | F |
| gp091 | - | 71,537 | 73,384 | tail fiber family protein | | structural components | F |
| gp092 | - | 73,455 | 73,742 | hypothetical protein | | | F |

| Locus_tag | Direction | Min | Max | Product | Notes | Function | Cluster |
|---|---|---|---|---|---|---|---|
| gp093 | - | 73,754 | 76,339 | tail spike protein | contains tail spike TSP1/Gp66 receptor binding N-terminal domain domain, and pectate lyase superfamily protein domain | structural components | F |

**Supplementary Table S5.** Genome annotations of *Klebsiella pneumoniae* phage vB_KpM_FBKp34.

| Locus_tag | Direction | Min | Max | Product | Notes | Function | Cluster |
|---|---|---|---|---|---|---|---|
| gp001 | + | 44 | 238 | hypothetical protein | | | D |
| gp002 | + | 344 | 1,012 | putative recombinase, resolvase family protein | contains resolvase, N terminal domain; similar to Phi92_gp052 | DNA recombination | D |
| gp003 | + | 1,020 | 1,232 | hypothetical protein | | | D |
| gp004 | + | 1,309 | 1,554 | hypothetical protein | | | D |
| gp005 | + | 1,616 | 1,948 | hypothetical protein | contains domain of unknown function DUF4326 | | D |
| gp006 | + | 1,948 | 2,232 | hypothetical protein | | | D |
| gp007 | + | 2,225 | 2,470 | hypothetical protein | | | D |
| gp008 | + | 2,470 | 2,718 | hypothetical protein | contains domain of unknown function DUF4236 | | D |
| gp009 | + | 2,727 | 2,819 | hypothetical protein | | | D |
| gp010 | + | 2,932 | 3,402 | hypothetical protein | | | D |
| gp011 | + | 3,406 | 3,729 | hypothetical protein | | | D |
| gp012 | + | 3,726 | 4,265 | hypothetical protein | contains domain of unknown function DUF1768 | | D |
| gp013 | + | 4,279 | 4,458 | hypothetical protein | | | D |
| gp014 | + | 4,593 | 4,805 | hypothetical protein | | | D |
| gp015 | + | 4,802 | 5,152 | hypothetical protein | | | D |
| gp016 | + | 5,149 | 5,370 | hypothetical protein | | | D |
| gp017 | + | 5,398 | 5,973 | hypothetical protein | | | D |
| gp018 | + | 5,966 | 6,415 | hypothetical protein | | | D |
| gp019 | + | 6,408 | 6,695 | hypothetical protein | | | D |
| gp020 | + | 6,685 | 7,011 | hypothetical protein | | | D |
| gp021 | + | 6,998 | 7,525 | hypothetical protein | | | D |
| gp022 | + | 7,522 | 7,764 | hypothetical protein | | | D |
| gp023 | + | 7,773 | 8,141 | hypothetical protein | | | D |

| **Locus_tag** | **Direction** | **Min** | **Max** | **Product** | **Notes** | **Function** | **Cluster** |
|---|---|---|---|---|---|---|---|
| gp024 | + | 8,143 | 8,370 | hypothetical protein | | | D |
| gp025 | + | 8,376 | 8,687 | hypothetical protein | | | D |
| gp026 | + | 8,695 | 8,943 | hypothetical protein | | | D |
| gp027 | + | 8,940 | 9,362 | hypothetical protein | | | D |
| gp028 | + | 9,369 | 9,548 | hypothetical protein | | | D |
| gp029 | + | 9,558 | 9,797 | hypothetical protein | contains restriction alleviation protein Lar domain | | D |
| gp030 | + | 9,863 | 10,240 | hypothetical protein | | | D |
| gp031 | + | 10,331 | 10,585 | hypothetical protein | | | D |
| gp032 | + | 10,582 | 10,863 | hypothetical protein | | | D |
| gp033 | + | 11,052 | 11,312 | hypothetical protein | | | D |
| gp034 | + | 11,322 | 11,543 | hypothetical protein | | | D |
| gp035 | + | 11,540 | 11,974 | hypothetical protein | | | D |
| gp036 | + | 11,977 | 12,144 | hypothetical protein | | | D |
| gp037 | + | 12,160 | 12,567 | hypothetical protein | | | D |
| gp038 | - | 13,697 | 13,855 | hypothetical protein | | | A |
| gp039 | - | 14,339 | 14,707 | hypothetical protein | contains phage GP30.8 protein domain | | A |
| gp040 | - | 14,776 | 15,144 | hypothetical protein | | | A |
| gp041 | - | 15,605 | 15,892 | hypothetical protein | | | A |
| gp042 | - | 15,903 | 16,088 | hypothetical protein | | | A |
| gp043 | - | 16,134 | 16,493 | hypothetical protein | | | A |
| gp044 | - | 16,579 | 16,962 | hypothetical protein | | | A |
| gp045 | - | 16,977 | 17,111 | hypothetical protein | | | A |
| gp046 | - | 17,185 | 17,535 | hypothetical protein | | | A |
| gp047 | - | 17,539 | 17,832 | hypothetical protein | | | A |
| gp048 | - | 17,878 | 18,177 | hypothetical protein | | | A |

| Locus_tag | Direction | Min | Max | Product | Notes | Function | Cluster |
|---|---|---|---|---|---|---|---|
| gp049 | - | 18,268 | 18,516 | hypothetical protein | | | A |
| gp050 | - | 18,578 | 18,802 | hypothetical protein | | | A |
| gp051 | - | 18,892 | 19,350 | hypothetical protein | | | A |
| gp052 | - | 19,689 | 19,850 | hypothetical protein | | | A |
| gp053 | - | 19,847 | 20,149 | hypothetical protein | | | A |
| gp054 | - | 20,149 | 20,409 | hypothetical protein | | | A |
| gp055 | - | 20,543 | 20,674 | hypothetical protein | | | A |
| gp056 | - | 20,677 | 20,967 | hypothetical protein | | | A |
| gp057 | - | 20,985 | 21,377 | hypothetical protein | | | A |
| gp058 | - | 21,456 | 21,800 | hypothetical protein | | | A |
| gp059 | - | 21,804 | 21,953 | hypothetical protein | | | A |
| gp060 | - | 22,009 | 22,359 | hypothetical protein | | | A |
| gp061 | - | 22,449 | 22,706 | hypothetical protein | | | A |
| gp062 | - | 22,752 | 22,973 | hypothetical protein | | | A |
| gp063 | - | 23,049 | 23,213 | hypothetical protein | | | A |
| gp064 | - | 23,291 | 23,533 | hypothetical protein | | | A |
| gp065 | - | 23,597 | 24,133 | hypothetical protein | | | A |
| gp066 | - | 24,223 | 24,444 | hypothetical protein | | | A |
| gp067 | - | 25,705 | 25,818 | putative DNA methylase | | DNA methylation | A |
| gp068 | - | 25,812 | 26,318 | HNH endonuclease | contains HNH endonuclease domain | DNA metabolism | A |
| gp069 | - | 26,324 | 27,016 | DNA cytosine methyltransferase | | DNA methylation | A |
| gp070 | - | 27,013 | 27,360 | hypothetical protein | | | A |
| gp071 | - | 27,350 | 27,751 | hypothetical protein | | | A |
| gp072 | - | 27,752 | 27,997 | hypothetical protein | | | A |

| Locus_tag | Direction | Min | Max | Product | Notes | Function | Cluster |
|---|---|---|---|---|---|---|---|
| gp073 | - | 28,001 | 29,545 | putative DNA helicase | contains AAA domain and UvrD-like helicase C-terminal domain | DNA replication | A |
| gp074 | - | 29,542 | 29,949 | hypothetical protein | | | A |
| gp075 | - | 29,946 | 30,263 | hypothetical protein | | | A |
| gp076 | - | 30,379 | 30,888 | hypothetical protein | | | A |
| gp077 | - | 30,899 | 32,182 | hypothetical protein | | | A |
| gp078 | - | 32,378 | 32,614 | hypothetical protein | contains restriction alleviation protein Lar domain | | A |
| gp079 | - | 32,752 | 34,104 | hypothetical protein | | | A |
| gp080 | - | 34,117 | 34,437 | hypothetical protein | | | A |
| gp081 | - | 34,444 | 34,692 | hypothetical protein | | | A |
| gp082 | - | 34,701 | 34,895 | hypothetical protein | | | A |
| gp083 | - | 34,892 | 35,089 | hypothetical protein | | | A |
| gp084 | - | 35,086 | 35,241 | hypothetical protein | | | A |
| gp085 | - | 35,241 | 35,336 | hypothetical protein | | | A |
| gp086 | - | 35,329 | 35,550 | hypothetical protein | | | A |
| gp087 | - | 35,550 | 35,918 | hypothetical protein | | | A |
| gp088 | - | 35,946 | 36,413 | hypothetical protein | | | A |
| gp089 | - | 36,424 | 36,558 | hypothetical protein | | | A |
| gp090 | - | 36,571 | 36,858 | hypothetical protein | | | A |
| gp091 | - | 36,842 | 37,396 | putative metal dependent phosphohydrolase | | | A |
| gp092 | + | 37,516 | 37,878 | hypothetical protein | | | B |
| gp093 | + | 37,881 | 38,252 | hypothetical protein | | | B |
| gp094 | + | 38,326 | 38,478 | hypothetical protein | | | B |
| gp095 | + | 38,478 | 38,801 | hypothetical protein | | | B |

| Locus_tag | Direction | Min | Max | Product | Notes | Function | Cluster |
|---|---|---|---|---|---|---|---|
| gp096 | + | 38,798 | 39,007 | hypothetical protein | | | B |
| gp097 | + | 39,000 | 39,341 | hypothetical protein | | | B |
| gp098 | + | 39,351 | 39,587 | hypothetical protein | | | B |
| gp099 | + | 39,590 | 40,288 | hypothetical protein | | | B |
| gp100 | + | 40,281 | 40,475 | hypothetical protein | contains RNA repair pathway DNA polymerase beta family domain | | B |
| gp101 | + | 40,476 | 41,237 | putative thioredoxin | | DNA metabolism | B |
| gp102 | + | 41,234 | 41,515 | hypothetical protein | | | B |
| gp103 | + | 41,699 | 42,076 | hypothetical protein | | | B |
| gp104 | + | 42,091 | 42,753 | hypothetical protein | | | B |
| gp105 | + | 42,762 | 43,277 | putative phosphatase | | | B |
| gp106 | + | 43,287 | 44,216 | anti-sigma factor | | | B |
| gp107 | + | 44,213 | 44,332 | hypothetical protein | | | B |
| gp108 | + | 44,336 | 44,746 | hypothetical protein | | | B |
| gp109 | + | 44,759 | 45,196 | hypothetical protein | | | B |
| gp110 | + | 45,224 | 45,661 | hypothetical protein | | | B |
| gp111 | + | 45,699 | 46,985 | poly [ADP-ribose] polymerase | contains WGR domain; Poly(ADP-ribose) polymerase, regulatory domain; and Poly(ADP-ribose) polymerase catalytic domain | DNA repair | B |
| gp112 | + | 47,003 | 47,470 | hypothetical protein | | | B |
| gp113 | + | 47,483 | 47,890 | hypothetical protein | contains Yqey-like protein domain | | B |
| gp114 | + | 47,975 | 50,992 | ribonucleotide reductase of class III (anaerobic), large subunit | contains ATP cone domain and anaerobic ribonucleoside-triphosphate reductase domains | DNA metabolism | B |
| gp115 | + | 51,002 | 51,253 | hypothetical protein | | | B |

| Locus_tag | Direction | Min | Max | Product | Notes | Function | Cluster |
|---|---|---|---|---|---|---|---|
| gp116 | + | 51,253 | 51,732 | ribonucleotide reductase of class III (anaerobic), activating protein | contains 4Fe-4S single cluster domain | DNA metabolism | B |
| gp117 | + | 51,713 | 52,096 | hypothetical protein | | | B |
| gp118 | + | 52,106 | 52,288 | hypothetical protein | | | B |
| gp119 | + | 52,281 | 52,703 | NinX | contains DUF2591 domain | | B |
| gp120 | + | 52,711 | 52,929 | hypothetical protein | | | B |
| gp121 | + | 52,926 | 53,258 | hypothetical protein | | | B |
| gp122 | + | 53,267 | 53,458 | bacterioferritin-associated ferredoxin | contains BFD-like [2Fe-2S] binding domain | | B |
| gp123 | + | 53,466 | 53,624 | hypothetical protein | | | B |
| gp124 | + | 53,653 | 54,630 | integral membrane protein TerC | contains integral membrane protein TerC family domain | | B |
| gp125 | + | 54,675 | 55,037 | hypothetical protein | | | B |
| gp126 | + | 55,180 | 55,353 | hypothetical protein | | | B |
| gp127 | + | 55,343 | 55,528 | hypothetical protein | | | B |
| gp128 | + | 55,529 | 55,732 | hypothetical protein | | | B |
| gp129 | + | 55,719 | 55,976 | hypothetical protein | | | B |
| gp130 | + | 55,976 | 56,308 | hypothetical protein | | | B |
| gp131 | + | 56,329 | 57,042 | putative tellurium resistance protein | contains vWA found in TerF C terminus domain | | B |
| gp132 | + | 57,110 | 57,691 | TerD family protein | contains TerD domain | | B |
| gp133 | + | 57,753 | 58,334 | TerD family protein | contains TerD domain | | B |
| gp134 | + | 58,393 | 59,271 | hypothetical protein | | | B |
| gp135 | + | 59,282 | 60,358 | toxic anion (tellurite) resistance protein | contains toxic anion resistance protein (TelA) domain | | B |
| gp136 | + | 60,423 | 60,932 | cell wall hydrolase | contains cell wall hydrolase domain | | B |
| gp137 | + | 60,945 | 61,196 | hypothetical protein | | | B |
| gp138 | + | 61,193 | 61,783 | hypothetical protein | | | B |
| gp139 | + | 61,815 | 62,117 | hypothetical protein | | | B |

| Locus_tag | Direction | Min | Max | Product | Notes | Function | Cluster |
|---|---|---|---|---|---|---|---|
| gp140 | + | 62,120 | 62,395 | hypothetical protein | | | B |
| gp141 | + | 62,395 | 63,045 | hypothetical protein | putative hydrolase | | B |
| gp142 | + | 63,045 | 63,869 | hypothetical protein | contains PRTase ComF-like domain | | B |
| gp143 | + | 63,879 | 64,298 | hypothetical protein | | | B |
| gp144 | + | 64,291 | 64,551 | hypothetical protein | | | B |
| gp145 | + | 64,544 | 64,963 | hypothetical protein | | | B |
| gp146 | + | 64,960 | 65,139 | hypothetical protein | | | B |
| gp147 | + | 65,136 | 65,552 | putative Eaa1 | | | B |
| gp148 | + | 65,549 | 66,055 | hypothetical protein | | | B |
| gp149 | + | 66,030 | 66,677 | hypothetical protein | | | B |
| gp150 | + | 66,714 | 66,965 | hypothetical protein | | | B |
| gp151 | + | 66,973 | 67,671 | hypothetical protein | | | B |
| gp152 | + | 67,685 | 69,484 | DNA primase/helicase | contains Toprim-like domain and DnaB-like helicase C terminal domain | DNA replication | B |
| gp153 | + | 69,501 | 71,717 | phage DNA polymerase I | contains DNA polymerase family A domain | DNA replication | B |
| gp154 | + | 71,810 | 72,670 | hypothetical protein | | | B |
| gp155 | - | 72,832 | 76,431 | hypothetical protein | contains chaperone of endosialidase domain | | C |
| gp156 | - | 76,431 | 76,916 | hypothetical protein | | | C |
| gp157 | - | 76,946 | 77,272 | hypothetical protein | | | C |
| gp158 | - | 77,274 | 77,762 | hypothetical protein | possible tail fiber assembly protein | | C |
| gp159 | - | 77,764 | 78,792 | hypothetical protein | | | C |
| gp160 | - | 78,805 | 79,428 | hypothetical protein | | | C |
| gp161 | - | 79,438 | 80,922 | putative baseplate | contains baseplate J-like protein domain | structural components | C |
| gp162 | - | 80,915 | 81,196 | putative membrane protein | | | C |
| gp163 | - | 81,210 | 81,734 | hypothetical protein | | | C |

| Locus_tag | Direction | Min | Max | Product | Notes | Function | Cluster |
|---|---|---|---|---|---|---|---|
| gp164 | - | 81,737 | 82,390 | baseplate central spike | contains phage protein gp137 N-terminal domain | structural components | C |
| gp165 | - | 82,391 | 83,356 | hypothetical protein | | | C |
| gp166 | - | 83,366 | 83,704 | hypothetical protein | | | C |
| gp167 | - | 83,716 | 84,417 | hypothetical protein | | | C |
| gp168 | - | 84,422 | 86,365 | putative cell envelope integrity protein TolA | | | C |
| gp169 | - | 86,377 | 86,598 | hypothetical protein | | | C |
| gp170 | - | 86,619 | 87,110 | hypothetical protein | | | C |
| gp171 | - | 87,187 | 87,654 | hypothetical protein | | | C |
| gp172 | - | 87,697 | 89,064 | putative tail sheath protein | contains DUF3383 domain | structural components | C |
| gp173 | - | 89,082 | 89,657 | hypothetical protein | | | C |
| gp174 | - | 89,650 | 90,069 | hypothetical protein | | | C |
| gp175 | - | 90,115 | 90,585 | hypothetical protein | | | C |
| gp176 | - | 90,585 | 91,091 | hypothetical protein | | | C |
| gp177 | - | 91,094 | 91,504 | hypothetical protein | | | C |
| gp178 | - | 91,571 | 92,569 | major capsid protein | contains phage major capsid protein E domain | structural components | C |
| gp179 | - | 92,587 | 92,988 | putative head stabilization/decoration protein | | structural components | C |
| gp180 | - | 93,004 | 94,344 | hypothetical protein | possible scaffold protein | | C |
| gp181 | - | 94,344 | 94,820 | hypothetical protein | contains putative phage serine protease XkdF domain | | C |
| gp182 | - | 94,968 | 96,548 | hypothetical protein | | | C |
| gp183 | - | 96,686 | 98,230 | putative terminase large subunit | contains terminase-like family domain and terminaseRNaseH-like domain | DNA packaging | C |
| gp184 | - | 98,352 | 98,651 | hypothetical protein | | | C |
| tRNA | - | 98,756 | 98,833 | tRNA-Lys-TTT | | | C |

| Locus_tag | Direction | Min | Max | Product | Notes | Function | Cluster |
|---|---|---|---|---|---|---|---|
| gp185 | - | 98,841 | 98,957 | hypothetical protein | | | C |
| tRNA | - | 98,980 | 99,055 | tRNA-Arg-TCT | | | C |
| gp186 | - | 99,071 | 99,250 | hypothetical protein | possible lysin | | C |
| tRNA | - | 99,267 | 99,342 | tRNA-Gly-TCC | | | C |
| tRNA | - | 99,647 | 99,723 | tRNA-His-GTG | | | C |
| tRNA | - | 99,964 | 100,055 | tRNA-Leu-TAA | | | C |
| tRNA | - | 100,067 | 100,141 | tRNA-Ile2-CAT | | | C |
| gp187 | - | 100,446 | 100,868 | hypothetical protein | possible Rz spanin | | C |
| gp188 | - | 101,442 | 102,566 | IS200/IS605 family element transposase accessory protein TnpB | contains helix-turn-helix domain; probable transposase domain; and putative transposase DNA-binding domain | DNA integration | C |
| gp189 | - | 102,999 | 103,373 | hypothetical protein | | | C |
| gp190 | - | 103,459 | 104,154 | hypothetical protein | | | C |
| gp191 | - | 104,273 | 104,596 | hypothetical protein | | | C |
| tRNA | - | 104,784 | 104,857 | tRNA-Trp-CCA | | | C |
| tRNA | - | 104,866 | 104,941 | tRNA-Phe-GAA | | | C |
| tRNA | - | 104,947 | 105,021 | tRNA-Asn-GTT | | | C |
| gp192 | - | 105,022 | 105,216 | hypothetical protein | | | C |
| tRNA | - | 105,263 | 105,340 | tRNA-Pro-TGG | | | C |
| tRNA | - | 105,347 | 105,421 | tRNA-Cys-GCA | | | C |
| tRNA | - | 105,596 | 105,671 | tRNA-Ile-GAT | | | C |
| tRNA | - | 105,678 | 105,763 | tRNA-Ser-GCT | | | C |
| tRNA | - | 105,844 | 105,929 | tRNA-Ser-TGA | | | C |
| tRNA | - | 105,936 | 106,013 | tRNA-Leu-TAG | | | C |
| tRNA | - | 106,024 | 106,101 | tRNA-Glu-TTC | | | C |
| tRNA | - | 106,110 | 106,223 | tRNA-Asp-GTC | | | C |

| Locus_tag | Direction | Min | Max | Product | Notes | Function | Cluster |
|---|---|---|---|---|---|---|---|
| tRNA | - | 106,230 | 106,306 | tRNA-Asp-GTC | | | C |
| gp193 | - | 106,786 | 108,225 | hypothetical protein | contains DUF2828 domain | | C |
| gp194 | - | 108,308 | 108,550 | hypothetical protein | | | C |
| gp195 | - | 108,624 | 109,478 | hypothetical protein | | | C |
| gp196 | - | 109,568 | 111,253 | nicotinamide phosphoribosyltransferase | contains DUF5598 domain and nicotinate phosphoribosyltransferase (NAPRTase) family domain | | C |
| gp197 | - | 111,243 | 111,836 | HNH homing endonuclease | contains NUMOD4 motif and HNH endonuclease domain | DNA metabolism | C |
| gp198 | - | 111,802 | 112,647 | ribose-phosphate pyrophosphokinase | contains N-terminal domain of ribose phosphate pyrophosphokinase and a phosphoribosyl transferase domain | DNA metabolism | C |
| gp199 | - | 112,644 | 113,027 | hypothetical protein | | | C |
| gp200 | + | 113,366 | 113,662 | hypothetical protein | | | D |
| gp201 | + | 113,662 | 114,708 | ribosylnicotinamide kinase | contains cytidylyltransferase-like domain and AAA domain | | D |
| gp202 | + | 114,723 | 114,908 | hypothetical protein | | | D |
| gp203 | + | 114,905 | 115,603 | nicotinamide mononucleotide transport | contains nicotinamide mononucleotide transporter domain | | D |
| gp204 | + | 115,633 | 115,950 | hypothetical protein | | | D |
| gp205 | + | 115,950 | 116,231 | hypothetical protein | | | D |
| gp206 | + | 116,484 | 117,119 | hypothetical protein | | | D |
| gp207 | + | 117,116 | 117,469 | hypothetical protein | possible restriction nuclease | | D |
| gp208 | + | 117,469 | 117,960 | hypothetical protein | | | D |
| gp209 | + | 117,962 | 118,519 | hypothetical protein | possible metallophosphoesterase | | D |
| gp210 | + | 118,678 | 119,598 | RNA ligase | contains RNA ligase domain | | D |
| gp211 | + | 119,598 | 119,942 | hypothetical protein | | | D |
| gp212 | + | 119,952 | 120,377 | hypothetical protein | contains AAA domain | | D |

| Locus_tag | Direction | Min | Max | Product | Notes | Function | Cluster |
|---|---|---|---|---|---|---|---|
| gp213 | + | 120,386 | 120,598 | hypothetical protein | | | D |
| gp214 | + | 120,598 | 121,374 | NAD-dependent protein deacetylase of SIR2 family | contains Sir2 family domain | | D |
| gp215 | + | 121,379 | 121,564 | hypothetical protein | | | D |
| gp216 | + | 121,567 | 121,785 | hypothetical protein | | | D |
| gp217 | + | 121,794 | 122,012 | hypothetical protein | | | D |
| gp218 | + | 122,009 | 123,370 | DNA ligase | contains ATP dependent DNA ligase domain | DNA replication and repair | D |
| gp219 | + | 123,372 | 124,031 | hypothetical protein | contains phosphoribosyl-ATP pyrophosphohydrolase domain | | D |
| gp220 | + | 124,039 | 124,146 | hypothetical protein | | | D |
| gp221 | + | 124,140 | 124,412 | hypothetical protein | | | D |
| gp222 | + | 124,425 | 125,360 | DNA recombination-dependent growth factor RdgC | contains putative exonuclease RdgC domain | DNA repair | D |
| gp223 | + | 125,418 | 125,645 | hypothetical protein | | | D |
| gp224 | + | 125,720 | 126,757 | exonuclease | | DNA replication | D |
| gp225 | + | 126,853 | 127,386 | hypothetical protein | | | D |
| gp226 | + | 127,350 | 127,934 | packaging and recombination endonuclease VII | contains recombination endonuclease VII domain | DNA repair | D |
| gp227 | + | 127,921 | 128,124 | hypothetical protein | | | D |
| gp228 | + | 128,121 | 129,059 | exonuclease | contains RNase_H superfamily domain | DNA replication | D |
| gp229 | + | 129,097 | 129,474 | hypothetical protein | | | D |
| gp230 | + | 129,474 | 129,641 | hypothetical protein | | | D |
| gp231 | + | 129,635 | 130,213 | hypothetical protein | contains 5' nucleotidase, deoxy (Pyrimidine), cytosolic type C protein (NT5C) domain | | D |

| **Locus_tag** | **Direction** | **Min** | **Max** | **Product** | **Notes** | **Function** | **Cluster** |
|---|---|---|---|---|---|---|---|
| gp232 | + | 130,210 | 130,533 | hypothetical protein | | | D |
| gp233 | + | 130,534 | 131,505 | putative thymidylate synthase | contains thymidylate synthase complementing protein domain | DNA metabolism | D |
| gp234 | + | 131,577 | 132,047 | hypothetical protein | | | D |
| gp235 | + | 132,061 | 132,312 | hypothetical protein | | | D |
| gp236 | + | 132,332 | 134,629 | ribonucleoside-diphosphate reductase | contains ATP cone domain and ribonucleotide reductase, barrel domain | DNA replication | D |
| gp237 | + | 134,671 | 135,771 | ribonucleotide-diphosphate reductase subunit beta | contains ribonucleotide reductase, small chain domain | DNA replication | D |
| gp238 | + | 135,784 | 136,038 | glutaredoxin | contains glutaredoxin domain | DNA metabolism | D |
| gp239 | + | 136,047 | 136,187 | hypothetical protein | | | D |
| gp240 | + | 136,187 | 136,588 | hypothetical protein | | | D |
| gp241 | + | 136,598 | 137,092 | lysozyme | contains phage lysozyme domain | cell lysis | D |
| gp242 | + | 137,170 | 137,916 | PhoH-like protein | contains PhoH-like protein domain | | D |
| gp243 | + | 137,923 | 138,384 | hypothetical protein | | | D |
| gp244 | + | 138,422 | 139,336 | hypothetical protein | | | D |
| gp245 | + | 139,290 | 140,063 | serine/threonine-protein phosphatase | contains calcineurin-like phosphoesterase domain | DNA repair | D |
| gp246 | + | 140,063 | 140,329 | hypothetical protein | | | D |
| gp247 | + | 140,326 | 140,892 | hypothetical protein | | | D |
| gp248 | + | 141,029 | 141,376 | putative DNA N-6-adenine methyltransferase | | DNA methylation | D |

**Supplementary Table S6.** Genome annotations of *Klebsiella pneumoniae* phage vB_KpM_FBKp24.

| **Locus_tag** | **Direction** | **Min** | **Max** | **Product** | **Notes** | **Function** |
|---|---|---|---|---|---|---|
| gp001 | - | 1 | 561 | hypothetical protein | | |
| gp002 | + | 619 | 2,178 | putative helicase | | DNA replication |
| gp003 | - | 2,262 | 2,627 | hypothetical protein | | |
| gp004 | - | 2,633 | 2,812 | hypothetical protein | | |
| gp005 | - | 2,895 | 3,533 | hypothetical protein | | |
| gp006 | - | 3,670 | 4,113 | hypothetical protein | N-acetyltransferase domain-containing protein | |
| gp007 | - | 4,221 | 5,564 | virion structural protein | | structural components |
| gp008 | - | 5,564 | 6,184 | hypothetical protein | | |
| gp009 | - | 6,184 | 7,557 | virion structural protein | | structural components |
| gp010 | - | 7,570 | 9,111 | hypothetical protein | | |
| gp011 | - | 9,180 | 10,103 | hypothetical protein | | |
| gp012 | - | 10,186 | 11,415 | hypothetical protein | | |
| gp013 | - | 11,425 | 12,666 | hypothetical protein | | |
| gp014 | - | 12,681 | 14,129 | hypothetical protein | | |
| gp015 | - | 14,221 | 16,647 | hypothetical protein | | |
| gp016 | - | 16,673 | 17,200 | putative virion structural protein | | structural components |
| gp017 | - | 17,206 | 18,111 | putative virion structural protein | | structural components |
| gp018 | - | 18,129 | 19,370 | virion structural protein | | structural components |
| gp019 | + | 19,407 | 20,582 | virion structural protein | | structural components |
| gp020 | + | 20,582 | 23,488 | virion structural protein | | structural components |
| gp021 | - | 23,532 | 24,764 | hypothetical protein | | |
| gp022 | - | 24,733 | 25,773 | hypothetical protein | | |
| gp023 | - | 25,784 | 26,242 | hypothetical protein | | |
| gp024 | - | 26,256 | 27,563 | putative virion structural protein | | structural components |

| Locus_tag | Direction | Min | Max | Product | Notes | Function |
|---|---|---|---|---|---|---|
| gp025 | + | 27,694 | 29,430 | putative DNA polymerase | | DNA replication |
| gp026 | - | 29,477 | 30,844 | putative DNA-directed RNA polymerase beta subunit | | transcription |
| gp027 | - | 30,837 | 31,676 | hypothetical protein | | |
| gp028 | + | 31,757 | 32,185 | hypothetical protein | | |
| gp029 | + | 32,178 | 33,893 | hypothetical protein | | |
| gp030 | + | 33,911 | 34,273 | hypothetical protein | | |
| gp031 | + | 34,288 | 34,926 | hypothetical protein | | |
| gp032 | - | 34,983 | 35,966 | hypothetical protein | | |
| gp033 | + | 36,038 | 38,032 | putative SNF2 domain/DEAD-like helicase | | transcription |
| gp034 | - | 38,094 | 38,477 | hypothetical protein | | |
| gp035 | - | 38,508 | 39,044 | hypothetical protein | | |
| gp036 | - | 39,126 | 39,314 | hypothetical protein | | |
| gp037 | - | 39,358 | 39,627 | hypothetical protein | possible DNA primase | |
| gp038 | - | 39,828 | 40,016 | hypothetical protein | | |
| gp039 | - | 40,013 | 40,489 | GNAT family N-acetyltransferase | | |
| gp040 | - | 40,683 | 40,835 | hypothetical protein | | |
| tRNA | - | 41,672 | 41,761 | tRNA-Ser-GCT | | |
| tRNA | - | 41,771 | 41,844 | tRNA-Asn-GTT | | |
| tRNA | - | 41,851 | 41,925 | tRNA-Asp-GTC | | |
| tRNA | - | 42,043 | 42,116 | tRNA-Gln-TTG | | |
| tRNA | - | 42,125 | 42,208 | tRNA-Tyr-GTA | | |
| gp041 | - | 42,392 | 42,751 | hypothetical protein | | |
| tRNA | - | 42,834 | 42,907 | tRNA-Arg-TCT | | |
| tRNA | - | 42,913 | 42,997 | tRNA-Leu-TAA | | |
| tRNA | - | 43,003 | 43,078 | tRNA-Ile2-CAT | | |

| Locus_tag | Direction | Min | Max | Product | Notes | Function |
|---|---|---|---|---|---|---|
| gp042 | - | 43,130 | 43,294 | hypothetical protein | | |
| gp043 | - | 43,298 | 44,155 | TRAP transporter solute receptor, TAXI family | contains a NMT1-like family domain | |
| gp044 | + | 44,395 | 44,652 | hypothetical protein | | |
| gp045 | - | 44,667 | 44,837 | hypothetical protein | | |
| gp046 | - | 44,849 | 45,736 | hypothetical protein | | |
| gp047 | - | 45,806 | 45,922 | hypothetical protein | | |
| tRNA | - | 46,041 | 46,120 | tRNA-Sup-TTA | | |
| gp048 | - | 46,370 | 47,137 | hypothetical protein | | |
| gp049 | - | 47,244 | 47,825 | hypothetical protein | | |
| gp050 | - | 47,883 | 48,305 | hypothetical protein | | |
| gp051 | - | 48,431 | 48,721 | hypothetical protein | | |
| gp052 | - | 48,733 | 49,902 | hypothetical protein | AAA domain-containing protein | |
| gp053 | - | 50,010 | 50,351 | hypothetical protein | | |
| gp054 | - | 50,438 | 51,706 | hypothetical protein | AAA domain-containing protein, putative ATP-binding protein | |
| gp055 | - | 51,735 | 52,370 | hypothetical protein | | |
| gp056 | - | 52,386 | 52,757 | hypothetical protein | putative tRNA nuclease WapA | |
| gp057 | - | 52,768 | 53,118 | hypothetical protein | | |
| gp058 | - | 53,151 | 53,510 | hypothetical protein | | |
| gp059 | - | 53,489 | 53,965 | hypothetical protein | | |
| gp060 | - | 53,976 | 54,578 | hypothetical protein | | |
| gp061 | - | 54,619 | 54,921 | hypothetical protein | | |
| gp062 | - | 55,283 | 56,239 | hypothetical protein | | |
| gp063 | - | 56,236 | 56,484 | hypothetical protein | | |
| gp064 | - | 56,510 | 56,875 | hypothetical protein | | |

| Locus_tag | Direction | Min | Max | Product | Notes | Function |
|---|---|---|---|---|---|---|
| gp065 | - | 56,891 | 57,337 | macro domain containing protein | | |
| gp066 | - | 57,815 | 58,279 | GNAT family N-acetyltransferase | | |
| gp067 | - | 58,503 | 60,017 | putative helicase | | DNA replication |
| gp068 | - | 60,173 | 62,140 | RNA polymerase beta subunit | putative ATP-dependent DNA helicase | transcription |
| gp069 | - | 62,140 | 64,413 | Putative DNA-directed RNA polymerase beta subunit | RNA_pol_Rpb2_6 domain-containing protein | transcription |
| gp070 | + | 64,527 | 64,877 | hypothetical protein | | |
| gp071 | - | 64,970 | 65,203 | hypothetical protein | | |
| gp072 | - | 65,263 | 66,489 | hypothetical protein | | |
| gp073 | - | 66,519 | 68,042 | hypothetical protein | | |
| gp074 | - | 68,070 | 69,611 | hypothetical protein | | |
| gp075 | - | 69,714 | 70,499 | hypothetical protein | | |
| gp076 | - | 70,570 | 71,415 | hypothetical protein | | |
| gp077 | - | 71,427 | 71,654 | hypothetical protein | | |
| gp078 | - | 71,657 | 72,778 | putative nuclease SbcCD D subunit | | DNA repair |
| gp079 | - | 72,768 | 73,325 | hypothetical protein | | |
| gp080 | - | 73,318 | 73,716 | hypothetical protein | | |
| gp081 | - | 73,809 | 74,429 | hypothetical protein | | |
| gp082 | - | 74,426 | 75,880 | Bifunctional DNA-directed RNA polymerase subunit beta-beta', RpoBC | | transcription |
| gp083 | - | 75,965 | 77,866 | hypothetical protein | low similarity (62% query cover, 24% ident, 2e-11 e-value) to nuclear shell protein gp104 from Pseudomonas phage 201phi2-1 | |
| gp084 | - | 79,189 | 81,405 | putative T4-like DNA polymerase | | DNA replication |
| gp085 | - | 81,517 | 81,939 | hypothetical protein | | |
| gp086 | - | 82,242 | 82,670 | hypothetical protein | | |

| Locus_tag | Direction | Min | Max | Product | Notes | Function |
|---|---|---|---|---|---|---|
| gp087 | - | 82,664 | 83,128 | hypothetical protein | | |
| gp088 | - | 83,137 | 83,562 | hypothetical protein | | |
| gp089 | - | 83,543 | 83,794 | hypothetical protein | | |
| gp090 | - | 83,907 | 84,182 | hypothetical protein | | |
| gp091 | - | 84,250 | 84,753 | hypothetical protein | | |
| gp092 | - | 84,765 | 85,169 | hypothetical protein | | |
| gp093 | - | 85,484 | 85,873 | hypothetical protein | | |
| gp094 | - | 86,120 | 87,088 | putative tubulin like protein | | |
| gp095 | + | 87,174 | 88,004 | hypothetical protein | | |
| gp096 | - | 88,049 | 88,882 | hypothetical protein | | |
| gp097 | - | 88,911 | 89,369 | hypothetical protein | | |
| gp098 | - | 89,398 | 90,018 | hypothetical protein | | |
| gp099 | - | 89,999 | 90,526 | hypothetical protein | | |
| gp100 | - | 90,587 | 91,141 | hypothetical protein | | |
| gp101 | - | 91,160 | 91,633 | hypothetical protein | | |
| gp102 | - | 91,620 | 92,111 | hypothetical protein | | |
| gp103 | - | 92,140 | 92,481 | hypothetical protein | | |
| gp104 | - | 92,551 | 93,231 | putative lytic transglycosidase | | |
| gp105 | - | 93,326 | 93,976 | hypothetical protein | | |
| gp106 | - | 94,186 | 94,596 | hypothetical protein | | |
| gp107 | - | 94,692 | 95,057 | hypothetical protein | | |
| gp108 | - | 95,047 | 95,412 | hypothetical protein | | |
| gp109 | - | 95,458 | 95,847 | hypothetical protein | | |
| gp110 | - | 95,888 | 96,208 | hypothetical protein | | |
| gp111 | - | 96,270 | 96,593 | hypothetical protein | | |

| Locus_tag | Direction | Min | Max | Product | Notes | Function |
|---|---|---|---|---|---|---|
| gp112 | + | 96,728 | 97,972 | putative head structural protein | | structural components |
| gp113 | - | 98,019 | 99,065 | hypothetical protein | | |
| gp114 | - | 99,169 | 101,343 | terminase large subunit | | DNA packaging |
| gp115 | - | 101,432 | 103,141 | putative virion structural protein | | structural components |
| gp116 | - | 103,143 | 105,692 | putative virion structural protein | | structural components |
| gp117 | - | 105,710 | 106,648 | virion structural protein | | structural components |
| gp118 | + | 106,812 | 108,881 | putative tail sheath protein | | structural components |
| gp119 | + | 108,874 | 109,749 | putative virion structural protein | putative baseplate wedge protein | structural components |
| gp120 | + | 109,880 | 110,524 | lytic transglycosylase | | |
| gp121 | + | 110,535 | 110,648 | hypothetical protein | | |
| gp122 | - | 110,703 | 111,299 | hypothetical protein | | |
| gp123 | - | 111,296 | 111,799 | hypothetical protein | | |
| gp124 | - | 111,771 | 112,715 | putative phosphohydrolase | | |
| gp125 | - | 112,771 | 113,019 | hypothetical protein | | |
| gp126 | - | 113,016 | 113,366 | hypothetical protein | | |
| gp127 | - | 113,463 | 113,864 | hypothetical protein | | |
| gp128 | - | 114,075 | 114,479 | hypothetical protein | | |
| gp129 | - | 114,543 | 114,725 | hypothetical protein | | |
| gp130 | - | 114,844 | 115,314 | hypothetical protein | | |
| gp131 | - | 115,405 | 115,683 | hypothetical protein | | |
| gp132 | - | 115,716 | 116,054 | hypothetical protein | | |
| gp133 | - | 116,075 | 116,635 | hypothetical protein | | |
| gp134 | - | 116,632 | 116,946 | hypothetical protein | | |
| gp135 | - | 117,340 | 117,687 | hypothetical protein | | |
| gp136 | - | 117,771 | 117,887 | hypothetical protein | | |

| Locus_tag | Direction | Min | Max | Product | Notes | Function |
|---|---|---|---|---|---|---|
| gp137 | - | 117,887 | 118,057 | hypothetical protein | | |
| gp138 | - | 118,072 | 118,230 | hypothetical protein | | |
| gp139 | + | 118,389 | 118,928 | hypothetical protein | | |
| gp140 | - | 118,969 | 119,388 | hypothetical protein | | |
| gp141 | - | 119,460 | 119,951 | hypothetical protein | | |
| gp142 | - | 120,012 | 120,416 | hypothetical protein | | |
| gp143 | - | 120,400 | 120,771 | hypothetical protein | | |
| gp144 | - | 120,839 | 121,255 | hypothetical protein | | |
| gp145 | - | 121,397 | 121,660 | hypothetical protein | | |
| gp146 | - | 121,743 | 122,186 | hypothetical protein | | |
| gp147 | - | 122,192 | 122,383 | hypothetical protein | | |
| gp148 | - | 122,383 | 122,715 | hypothetical protein | | |
| gp149 | - | 122,699 | 122,920 | hypothetical protein | | |
| gp150 | - | 122,926 | 123,333 | hypothetical protein | | |
| gp151 | - | 123,330 | 123,827 | hypothetical protein | | |
| gp152 | - | 123,796 | 124,494 | guanosine-3',5'-bis(diphosphate) 3'-pyrophosphohydrolase / GTP pyrophosphokinase, (p)ppGpp synthetase II | | regulation |
| gp153 | - | 124,602 | 125,225 | hypothetical protein | | |
| gp154 | - | 125,281 | 126,555 | hypothetical protein | | |
| gp155 | + | 126,744 | 128,039 | hypothetical protein | | |
| gp156 | - | 128,095 | 128,460 | hypothetical protein | | |
| gp157 | - | 128,482 | 128,835 | hypothetical protein | | |
| gp158 | - | 128,838 | 129,581 | hypothetical protein | | |
| gp159 | - | 129,589 | 130,161 | hypothetical protein | | |
| gp160 | - | 130,163 | 130,426 | hypothetical protein | | |

| Locus_tag | Direction | Min | Max | Product | Notes | Function |
|---|---|---|---|---|---|---|
| gp161 | - | 130,428 | 131,018 | thymidine kinase | | DNA metabolism |
| gp162 | - | 131,020 | 131,295 | hypothetical protein | | |
| gp163 | - | 131,292 | 131,651 | hypothetical protein | | |
| gp164 | - | 131,705 | 132,310 | hypothetical protein | | |
| gp165 | - | 132,315 | 132,683 | hypothetical protein | | |
| gp166 | - | 132,680 | 133,267 | hypothetical protein | | |
| gp167 | - | 133,281 | 133,568 | hypothetical protein | | |
| gp168 | + | 133,771 | 136,416 | hypothetical protein | | |
| gp169 | - | 136,450 | 136,710 | hypothetical protein | | |
| gp170 | - | 136,707 | 137,111 | hypothetical protein | | |
| gp171 | - | 137,166 | 137,978 | phosphate starvation-inducible protein PhoH, predicted ATPase | | |
| gp172 | - | 138,073 | 138,420 | hypothetical protein | | |
| gp173 | - | 138,499 | 138,861 | hypothetical protein | | |
| gp174 | - | 138,874 | 139,173 | hypothetical protein | | |
| gp175 | - | 139,225 | 139,692 | hypothetical protein | | |
| gp176 | + | 139,828 | 141,288 | glycosyltransferase | | |
| gp177 | + | 141,307 | 141,897 | hypothetical protein | | |
| gp178 | - | 141,959 | 142,372 | hypothetical protein | | |
| gp179 | - | 142,694 | 143,161 | hypothetical protein | | |
| gp180 | - | 143,180 | 143,707 | DUF1190 domain containing protein | | |
| gp181 | - | 143,786 | 144,178 | hypothetical protein | | |
| gp182 | - | 144,175 | 144,624 | hypothetical protein | | |
| gp183 | - | 144,677 | 144,976 | hypothetical protein | | |
| gp184 | - | 144,969 | 145,424 | hypothetical protein | | |

| Locus_tag | Direction | Min | Max | Product | Notes | Function |
|---|---|---|---|---|---|---|
| gp185 | - | 145,421 | 145,873 | hypothetical protein | | |
| gp186 | - | 145,875 | 146,459 | DUF1653 domain-containing protein | | |
| gp187 | - | 146,532 | 147,173 | hypothetical protein | | |
| gp188 | - | 147,177 | 147,596 | hypothetical protein | | |
| gp189 | - | 147,627 | 148,154 | hypothetical protein | | |
| gp190 | - | 148,199 | 149,713 | hypothetical protein | | |
| gp191 | - | 149,772 | 150,050 | hypothetical protein | | |
| gp192 | - | 150,082 | 150,450 | hypothetical protein | | |
| gp193 | - | 150,591 | 152,348 | nicotinamide phosphoribosyltransferase | | |
| gp194 | - | 152,345 | 153,208 | ribose-phosphate pyrophosphokinase | | |
| gp195 | - | 153,257 | 153,796 | dihydrofolate reductase | | |
| gp196 | + | 153,978 | 155,960 | putative tail fiber protein | contains head binding domain of P22 tailspike | structural components |
| gp197 | - | 156,038 | 156,760 | HigA family addiction module antidote protein | contains helix-turn-helix domain | |
| gp198 | - | 156,816 | 157,229 | hypothetical protein | | |
| gp199 | - | 157,271 | 157,666 | hypothetical protein | | |
| gp200 | - | 157,668 | 158,078 | hypothetical protein | | |
| gp201 | - | 158,091 | 158,456 | hypothetical protein | | |
| gp202 | - | 158,588 | 159,394 | hypothetical protein | | |
| gp203 | - | 159,510 | 160,040 | hypothetical protein | | |
| gp204 | - | 160,059 | 160,499 | hypothetical protein | | |
| gp205 | - | 160,496 | 160,876 | hypothetical protein | | |
| gp206 | - | 160,942 | 162,630 | hypothetical protein | | |
| gp207 | - | 162,685 | 163,278 | hypothetical protein | | |
| gp208 | - | 163,268 | 163,636 | hypothetical protein | | |
| gp209 | - | 163,640 | 163,969 | hypothetical protein | | |

| Locus_tag | Direction | Min | Max | Product | Notes | Function |
|---|---|---|---|---|---|---|
| gp210 | - | 164,007 | 164,783 | hypothetical protein | | |
| gp211 | - | 164,811 | 165,143 | hypothetical protein | | |
| gp212 | - | 165,240 | 165,713 | hypothetical protein | | |
| gp213 | - | 165,732 | 166,067 | hypothetical protein | | |
| gp214 | - | 166,079 | 166,405 | hypothetical protein | | |
| gp215 | - | 166,406 | 166,630 | hypothetical protein | | |
| gp216 | - | 166,620 | 167,009 | hypothetical protein | | |
| gp217 | - | 167,012 | 167,455 | hypothetical protein | | |
| gp218 | - | 167,455 | 168,429 | hypothetical protein | | |
| gp219 | - | 168,416 | 168,979 | hypothetical protein | | |
| gp220 | - | 169,006 | 169,194 | hypothetical protein | | |
| gp221 | - | 169,211 | 169,486 | hypothetical protein | | |
| gp222 | - | 169,486 | 169,746 | hypothetical protein | | |
| gp223 | - | 169,756 | 170,079 | hypothetical protein | | |
| gp224 | - | 170,082 | 170,588 | hypothetical protein | | |
| gp225 | - | 170,725 | 172,029 | AAA family ATPase | | |
| gp226 | - | 172,147 | 172,497 | hypothetical protein | | |
| gp227 | - | 172,494 | 172,949 | hypothetical protein | | |
| gp228 | - | 172,949 | 173,257 | hypothetical protein | | |
| gp229 | - | 173,261 | 174,952 | hypothetical protein | | |
| gp230 | - | 174,955 | 176,199 | putative tRNA-splicing ligase RtcB | | RNA repair |
| gp231 | - | 176,214 | 176,933 | hypothetical protein | | |
| gp232 | - | 176,952 | 177,515 | hypothetical protein | | |
| gp233 | - | 177,518 | 177,787 | hypothetical protein | | |
| gp234 | - | 177,777 | 178,289 | hypothetical protein | | |

| Locus_tag | Direction | Min | Max | Product | Notes | Function |
|---|---|---|---|---|---|---|
| gp235 | - | 178,289 | 178,663 | hypothetical protein | | |
| gp236 | - | 178,665 | 179,048 | hypothetical protein | | |
| gp237 | - | 179,045 | 179,290 | hypothetical protein | | |
| gp238 | + | 179,325 | 179,657 | putative ammonia monooxygenase | | |
| gp239 | - | 179,705 | 181,096 | AAA family ATPase | | |
| gp240 | - | 181,194 | 181,358 | hypothetical protein | | |
| gp241 | - | 181,418 | 181,801 | hypothetical protein | | |
| gp242 | - | 181,802 | 182,167 | hypothetical protein | | |
| gp243 | - | 182,173 | 182,646 | hypothetical protein | macro domain containing protein | |
| gp244 | - | 182,665 | 183,282 | hypothetical protein | | |
| gp245 | - | 183,288 | 183,887 | hypothetical protein | | |
| gp246 | - | 183,897 | 184,304 | hypothetical protein | | |
| gp247 | - | 184,308 | 185,525 | hypothetical protein | | |
| gp248 | - | 185,585 | 185,893 | hypothetical protein | | |
| gp249 | - | 185,893 | 186,024 | hypothetical protein | | |
| gp250 | - | 186,021 | 187,412 | thymidylate synthase | | DNA metabolism |
| gp251 | - | 187,409 | 187,642 | hypothetical protein | | |
| gp252 | - | 187,636 | 188,031 | hypothetical protein | | |
| gp253 | - | 188,021 | 188,536 | hypothetical protein | | |
| gp254 | - | 188,610 | 189,209 | tail protein | | structural components |
| gp255 | - | 189,214 | 189,699 | hypothetical protein | | |
| gp256 | - | 189,723 | 190,133 | hypothetical protein | | |
| gp257 | - | 190,193 | 190,555 | hypothetical protein | | |
| gp258 | - | 190,552 | 191,109 | hypothetical protein | | |
| gp259 | - | 191,106 | 191,669 | RNA 2'-phosphotransferase | | |

| Locus_tag | Direction | Min | Max | Product | Notes | Function |
|---|---|---|---|---|---|---|
| gp260 | - | 191,662 | 192,180 | hypothetical protein | | |
| gp261 | - | 192,180 | 192,455 | hypothetical protein | | |
| gp262 | - | 192,442 | 193,161 | hypothetical protein | | |
| gp263 | - | 193,154 | 193,867 | hypothetical protein | | |
| gp264 | - | 193,867 | 194,355 | hypothetical protein | | |
| gp265 | - | 194,470 | 194,877 | hypothetical protein | | |
| gp266 | - | 194,958 | 195,506 | hypothetical protein | | |
| gp267 | - | 195,507 | 195,851 | hypothetical protein | | |
| gp268 | - | 195,916 | 197,055 | hypothetical protein | | |
| gp269 | - | 197,052 | 197,240 | hypothetical protein | | |
| gp270 | - | 197,261 | 197,665 | hypothetical protein | | |
| gp271 | - | 197,667 | 198,602 | hypothetical protein | | |
| gp272 | - | 198,613 | 198,960 | hypothetical protein | | |
| gp273 | - | 199,043 | 199,555 | hypothetical protein | | |
| gp274 | - | 199,572 | 200,021 | membrane protease subunit | | |
| gp275 | - | 200,021 | 200,272 | hypothetical protein | | |
| gp276 | - | 200,300 | 200,989 | nicotinamide mononucleotide transporter | | |
| gp277 | - | 200,986 | 201,138 | hypothetical protein | | |
| gp278 | - | 201,152 | 202,366 | multifunctional transcriptional regulator/nicotinamide-nucleotide adenylyltransferase/ribosylnicotinamide kinase NadR | | |
| gp279 | - | 202,386 | 202,697 | hypothetical protein | | |
| gp280 | - | 202,710 | 203,105 | hypothetical protein | | |
| gp281 | - | 203,114 | 203,347 | hypothetical protein | | |
| gp282 | - | 203,357 | 203,917 | hypothetical protein | | |

| Locus_tag | Direction | Min | Max | Product | Notes | Function |
|---|---|---|---|---|---|---|
| gp283 | - | 203,918 | 204,178 | hypothetical protein | | |
| gp284 | - | 204,189 | 204,314 | hypothetical protein | | |
| gp285 | - | 204,379 | 205,014 | hypothetical protein | | |
| gp286 | - | 205,011 | 205,304 | hypothetical protein | | |
| gp287 | - | 205,301 | 205,735 | hypothetical protein | | |
| gp288 | - | 205,760 | 206,251 | hypothetical protein | | |
| gp289 | - | 206,229 | 206,507 | hypothetical protein | | |
| gp290 | - | 206,509 | 206,736 | hypothetical protein | | |
| gp291 | - | 206,747 | 207,154 | hypothetical protein | | |
| gp292 | - | 207,142 | 207,570 | hypothetical protein | | |
| gp293 | - | 207,589 | 208,176 | deoxyuridine 5'-triphosphate nucleotidohydrolase | | DNA metabolism |
| gp294 | - | 208,276 | 210,345 | bifunctional tail protein | | structural components |
| gp295 | - | 210,361 | 212,400 | bifunctional tail protein | | structural components |
| gp296 | - | 212,532 | 212,933 | hypothetical protein | | |
| gp297 | - | 212,930 | 213,259 | hypothetical protein | | |
| gp298 | + | 213,383 | 213,595 | hypothetical protein | | |
| gp299 | - | 213,638 | 213,883 | hypothetical protein | | |
| gp300 | - | 214,060 | 215,856 | putative tail fiber protein | contains beta-helix domain | structural components |
| gp301 | - | 215,902 | 218,649 | hypothetical protein | tail spike N-terminal domain containing protein | |
| gp302 | - | 218,757 | 218,951 | hypothetical protein | | |
| gp303 | - | 218,951 | 220,936 | putative tail fiber protein | | structural components |
| gp304 | - | 221,011 | 223,278 | putative tail fiber protein | | structural components |
| gp305 | - | 223,346 | 223,639 | hypothetical protein | | |
| gp306 | - | 223,651 | 226,302 | putative tail fiber protein | | structural components |
| gp307 | - | 226,470 | 228,698 | hypothetical protein | beta-helix domain containing protein | |

| Locus_tag | Direction | Min | Max | Product | Notes | Function |
|---|---|---|---|---|---|---|
| gp308 | + | 228,908 | 231,121 | hypothetical protein | putative tail fiber or spike with GTPase (hydrolase) activity | |
| gp309 | + | 231,199 | 233,454 | hypothetical protein | exopolysaccharide biosynthesis protein; putative tail fiber | |
| gp310 | + | 233,520 | 235,559 | hypothetical protein | peptidase S74 domain-containing protein; putative tail fiber | |
| gp311 | + | 235,573 | 235,794 | hypothetical protein | | |
| gp312 | + | 235,874 | 237,868 | hypothetical protein | putative tail spike protein | |
| gp313 | + | 237,880 | 239,541 | hypothetical protein | | |
| gp314 | - | 239,593 | 240,159 | hypothetical protein | putative virion structural protein | |
| gp315 | - | 240,210 | 240,596 | hypothetical protein | | |
| gp316 | - | 240,660 | 241,184 | hypothetical protein | | |
| gp317 | - | 241,239 | 241,790 | hypothetical protein | | |
| gp318 | - | 241,883 | 242,344 | hypothetical protein | | |
| gp319 | - | 242,505 | 243,155 | hypothetical protein | | |
| gp320 | - | 243,227 | 243,958 | hypothetical protein | | |
| gp321 | - | 244,085 | 244,246 | hypothetical protein | | |
| gp322 | - | 244,285 | 244,974 | hypothetical protein | | |
| gp323 | - | 245,080 | 245,826 | virion structural protein | possible prohead core protein protease | structural components |
| gp324 | - | 245,830 | 246,558 | hypothetical protein | | |
| gp325 | - | 246,560 | 247,285 | hypothetical protein | | |
| gp326 | - | 247,297 | 248,829 | hypothetical protein | | |
| gp327 | - | 248,829 | 249,020 | hypothetical protein | | |
| gp328 | - | 249,101 | 253,579 | RNA polymerase beta subunit | | transcription |
| gp329 | - | 253,596 | 255,227 | putative DNA-directed RNA polymerase beta subunit | | transcription |

| Locus_tag | Direction | Min | Max | Product | Notes | Function |
|---|---|---|---|---|---|---|
| gp330 | + | 255,274 | 261,879 | hypothetical protein | putative tail fiber with transglycosylase activity | |
| gp331 | + | 261,933 | 264,077 | hypothetical protein | | |
| gp332 | - | 264,119 | 264,463 | hypothetical protein | | |
| gp333 | - | 264,547 | 265,614 | hypothetical protein | | |
| gp334 | - | 265,605 | 265,898 | hypothetical protein | | |
| gp335 | - | 265,980 | 266,735 | hypothetical protein | | |
| gp336 | - | 266,748 | 267,266 | hypothetical protein | | |
| gp337 | + | 267,395 | 267,916 | hypothetical protein | | |
| gp338 | - | 267,956 | 268,657 | hypothetical protein | | |
| gp339 | - | 268,654 | 269,130 | hypothetical protein | | |
| gp340 | - | 269,135 | 269,806 | virion structural protein | | structural components |
| gp341 | - | 270,182 | 270,469 | hypothetical protein | | |
| gp342 | - | 270,488 | 271,903 | UvsX DNA recombinase | | DNA replication and repair |
| gp343 | + | 272,012 | 272,884 | hypothetical protein | | |
| gp344 | - | 272,927 | 273,304 | hypothetical protein | | |
| gp345 | - | 273,301 | 274,899 | RnhA ribonuclease HI | | DNA replication |
| gp346 | - | 274,956 | 275,384 | hypothetical protein | | |
| gp347 | + | 275,443 | 276,744 | putative virion structural protein | | structural components |
| gp348 | + | 276,747 | 277,349 | hypothetical protein | | |
| gp349 | - | 277,424 | 277,675 | hypothetical protein | | |
| gp350 | - | 277,783 | 278,028 | hypothetical protein | | |
| gp351 | - | 278,030 | 278,353 | hypothetical protein | | |
| gp352 | - | 278,343 | 278,984 | hypothetical protein | | |
| gp353 | - | 279,077 | 281,548 | nuclease SbcCD subunit C | | DNA repair |

| Locus_tag | Direction | Min | Max | Product | Notes | Function |
|---|---|---|---|---|---|---|
| gp354 | + | 281,627 | 282,616 | hypothetical protein | | |
| gp355 | - | 282,660 | 283,190 | hypothetical protein | | |
| gp356 | - | 283,206 | 284,561 | YomR | | |
| gp357 | - | 284,575 | 286,806 | putative tail fiber protein | | structural components |
| gp358 | + | 286,872 | 288,116 | putative virion structural protein | | structural components |
| gp359 | + | 288,119 | 288,718 | putative spanin | | cell lysis |
| gp360 | - | 288,763 | 289,380 | hypothetical protein | | |
| gp361 | - | 289,380 | 289,997 | thymidylate kinase | | DNA metabolism |
| gp362 | - | 290,010 | 290,591 | hypothetical protein | possible RuvC-like resolvase | |
| gp363 | - | 290,677 | 291,822 | hypothetical protein | | |
| gp364 | - | 291,837 | 292,754 | virion structural protein | | structural components |
| gp365 | - | 292,864 | 293,406 | hypothetical protein | | |
| gp366 | - | 293,440 | 294,282 | putative virion structural protein | | structural components |
| gp367 | - | 294,362 | 296,806 | virion structural protein | | structural components |
| gp368 | + | 296,871 | 299,660 | virion structural protein | | structural components |
| gp369 | - | 299,732 | 301,936 | hypothetical protein | | |
| gp370 | - | 301,967 | 303,577 | hypothetical protein | putative RNA polymerase beta subunit | |
| gp371 | - | 303,599 | 304,594 | hypothetical protein | | |
| gp372 | - | 304,809 | 307,097 | major head protein | | structural components |

**Supplementary Table S7.** Analysis of codon usage of phages vB_KpP_FBKp27, vB_KpM_FBKp34, vB_KpM_FBKp24 and *Klebsiella pneumoniae* HS11286 using Cusp from EMBOSS.

| Codon | AA[a] | ϕKp27 | | | | ϕKp34 | | | | ϕKp24 | | | | *K. pneumoniae* HS11286 | | |
|---|---|---|---|---|---|---|---|---|---|---|---|---|---|---|---|---|
| | | Fraction[b] | Frequency | No. | Ratio to bacteria | Fraction | Frequency | No. | Ratio to bacteria | Fraction | Frequency | No. | Ratio to bacteria | Fraction | Frequency | No. |
| GCA | A | 0.25 | 23.25 | 549 | 2.93 | 0.43 | 25.15 | 1061 | 3.17 | 0.25 | 16.80 | 1597 | 2.12 | 0.08 | 7.94 | 12537 |
| GCC | A | 0.16 | 14.70 | 347 | 0.34 | 0.11 | 6.38 | 269 | 0.15 | 0.21 | 14.36 | 1365 | 0.33 | 0.42 | 43.25 | 68298 |
| GCG | A | 0.03 | 2.50 | 59 | 0.06 | 0.08 | 4.46 | 188 | 0.10 | 0.31 | 21.04 | 2000 | 0.49 | 0.41 | 42.99 | 67889 |
| GCT | A | 0.56 | 51.04 | 1205 | 5.06 | 0.38 | 22.07 | 931 | 2.19 | 0.24 | 16.26 | 1545 | 1.61 | 0.10 | 10.09 | 15928 |
| TGC | C | 0.34 | 3.18 | 75 | 0.40 | 0.25 | 3.34 | 141 | 0.42 | 0.42 | 3.63 | 345 | 0.46 | 0.73 | 7.98 | 12595 |
| TGT | C | 0.66 | 6.14 | 145 | 2.03 | 0.75 | 10.12 | 427 | 3.35 | 0.58 | 5.06 | 481 | 1.67 | 0.28 | 3.03 | 4777 |
| GAC | D | 0.28 | 16.60 | 392 | 0.70 | 0.23 | 14.53 | 613 | 0.61 | 0.45 | 26.95 | 2561 | 1.13 | 0.47 | 23.82 | 37612 |
| GAT | D | 0.72 | 43.63 | 1030 | 1.64 | 0.77 | 49.87 | 2104 | 1.88 | 0.56 | 33.62 | 3195 | 1.26 | 0.53 | 26.60 | 42007 |
| GAA | E | 0.64 | 40.92 | 966 | 1.49 | 0.72 | 53.59 | 2261 | 1.96 | 0.68 | 44.74 | 4252 | 1.63 | 0.51 | 27.39 | 43255 |
| GAG | E | 0.36 | 22.91 | 541 | 0.86 | 0.28 | 20.93 | 883 | 0.79 | 0.32 | 20.65 | 1963 | 0.77 | 0.49 | 26.66 | 42100 |
| TTC | F | 0.55 | 19.65 | 464 | 1.01 | 0.30 | 13.51 | 570 | 0.70 | 0.49 | 23.30 | 2214 | 1.20 | 0.51 | 19.42 | 30677 |
| TTT | F | 0.46 | 16.39 | 387 | 0.87 | 0.70 | 31.62 | 1334 | 1.68 | 0.51 | 23.92 | 2273 | 1.27 | 0.49 | 18.81 | 29712 |
| GGA | G | 0.10 | 6.27 | 148 | 1.06 | 0.25 | 14.84 | 626 | 2.52 | 0.17 | 10.66 | 1013 | 1.81 | 0.08 | 5.89 | 9306 |
| GGC | G | 0.19 | 12.41 | 293 | 0.29 | 0.11 | 6.87 | 290 | 0.16 | 0.21 | 13.09 | 1244 | 0.31 | 0.56 | 42.48 | 67081 |
| GGG | G | 0.14 | 9.06 | 214 | 0.63 | 0.11 | 6.80 | 287 | 0.47 | 0.25 | 16.11 | 1531 | 1.11 | 0.19 | 14.47 | 22847 |
| GGT | G | 0.57 | 36.98 | 873 | 2.84 | 0.53 | 31.91 | 1346 | 2.45 | 0.38 | 24.11 | 2291 | 1.85 | 0.17 | 13.00 | 20537 |
| CAC | H | 0.40 | 6.99 | 165 | 0.62 | 0.30 | 5.67 | 239 | 0.50 | 0.48 | 7.77 | 738 | 0.69 | 0.50 | 11.34 | 17901 |
| CAT | H | 0.60 | 10.46 | 247 | 0.92 | 0.71 | 13.54 | 571 | 1.19 | 0.52 | 8.55 | 813 | 0.75 | 0.50 | 11.41 | 18018 |
| ATA | I | 0.08 | 4.41 | 104 | 1.43 | 0.16 | 10.95 | 462 | 3.55 | 0.10 | 6.53 | 621 | 2.12 | 0.06 | 3.09 | 4877 |
| ATC | I | 0.45 | 25.03 | 591 | 0.78 | 0.24 | 16.40 | 692 | 0.51 | 0.43 | 27.66 | 2629 | 0.86 | 0.58 | 32.23 | 50902 |
| ATT | I | 0.47 | 26.05 | 615 | 1.28 | 0.60 | 40.30 | 1700 | 1.98 | 0.47 | 30.78 | 2925 | 1.51 | 0.37 | 20.34 | 32115 |
| AAA | K | 0.58 | 37.32 | 881 | 1.51 | 0.74 | 61.56 | 2597 | 2.49 | 0.73 | 41.80 | 3973 | 1.69 | 0.64 | 24.68 | 38981 |
| AAG | K | 0.42 | 27.02 | 638 | 1.96 | 0.26 | 21.50 | 907 | 1.56 | 0.28 | 15.88 | 1509 | 1.15 | 0.36 | 13.77 | 21748 |

| Codon | AA[a] | ϕKp27 | | | | ϕKp34 | | | | ϕKp24 | | | | *K. pneumoniae* HS11286 | | |
|---|---|---|---|---|---|---|---|---|---|---|---|---|---|---|---|---|
| | | Fraction[b] | Frequency | No. | Ratio to bacteria | Fraction | Frequency | No. | Ratio to bacteria | Fraction | Frequency | No. | Ratio to bacteria | Fraction | Frequency | No. |
| CTA | L | 0.06 | 4.83 | 114 | 1.62 | 0.09 | 7.70 | 325 | 2.58 | 0.06 | 5.52 | 525 | 1.85 | 0.03 | 2.98 | 4711 |
| CTC | L | 0.11 | 9.40 | 222 | 0.61 | 0.05 | 4.39 | 185 | 0.29 | 0.16 | 14.03 | 1333 | 0.91 | 0.14 | 15.35 | 24245 |
| CTG | L | 0.41 | 35.32 | 834 | 0.49 | 0.12 | 10.00 | 422 | 0.14 | 0.22 | 19.37 | 1841 | 0.27 | 0.65 | 71.38 | 112738 |
| CTT | L | 0.18 | 15.54 | 367 | 2.00 | 0.28 | 23.09 | 974 | 2.97 | 0.17 | 15.00 | 1426 | 1.93 | 0.07 | 7.79 | 12296 |
| TTA | L | 0.16 | 14.23 | 336 | 2.27 | 0.31 | 25.60 | 1080 | 4.08 | 0.24 | 21.55 | 2048 | 3.43 | 0.06 | 6.28 | 9910 |
| TTG | L | 0.09 | 7.50 | 177 | 1.11 | 0.15 | 12.40 | 523 | 1.84 | 0.16 | 13.95 | 1326 | 2.07 | 0.06 | 6.74 | 10649 |
| ATG | M | 1.00 | 29.10 | 687 | 1.09 | 1.00 | 24.34 | 1027 | 0.91 | 1.00 | 21.21 | 2016 | 0.79 | 1.00 | 26.77 | 42285 |
| AAC | N | 0.53 | 27.95 | 660 | 1.27 | 0.30 | 16.14 | 681 | 0.73 | 0.53 | 28.19 | 2679 | 1.28 | 0.65 | 21.98 | 34709 |
| AAT | N | 0.47 | 24.65 | 582 | 2.04 | 0.70 | 37.26 | 1572 | 3.08 | 0.47 | 25.12 | 2387 | 2.08 | 0.36 | 12.09 | 19101 |
| CCA | P | 0.31 | 12.20 | 288 | 3.05 | 0.41 | 12.14 | 512 | 3.04 | 0.19 | 7.50 | 713 | 1.88 | 0.09 | 4.00 | 6310 |
| CCC | P | 0.10 | 3.73 | 88 | 0.49 | 0.05 | 1.54 | 65 | 0.20 | 0.15 | 5.87 | 558 | 0.77 | 0.16 | 7.59 | 11987 |
| CCG | P | 0.09 | 3.60 | 85 | 0.12 | 0.09 | 2.77 | 117 | 0.09 | 0.37 | 14.55 | 1383 | 0.48 | 0.65 | 30.15 | 47615 |
| CCT | P | 0.50 | 19.61 | 463 | 4.18 | 0.45 | 13.39 | 565 | 2.86 | 0.29 | 11.65 | 1107 | 2.48 | 0.10 | 4.69 | 7406 |
| CAA | Q | 0.38 | 17.41 | 411 | 2.52 | 0.68 | 22.02 | 929 | 3.19 | 0.54 | 17.69 | 1681 | 2.56 | 0.15 | 6.91 | 10906 |
| CAG | Q | 0.62 | 28.67 | 677 | 0.75 | 0.32 | 10.31 | 435 | 0.27 | 0.46 | 15.32 | 1456 | 0.40 | 0.85 | 38.05 | 60087 |
| AGA | R | 0.10 | 4.36 | 103 | 2.62 | 0.35 | 13.44 | 567 | 8.06 | 0.12 | 6.02 | 572 | 3.61 | 0.03 | 1.67 | 2632 |
| AGG | R | 0.06 | 2.41 | 57 | 1.45 | 0.10 | 3.67 | 155 | 2.20 | 0.05 | 2.75 | 261 | 1.65 | 0.03 | 1.67 | 2636 |
| CGA | R | 0.08 | 3.56 | 84 | 1.35 | 0.09 | 3.56 | 150 | 1.35 | 0.13 | 6.82 | 648 | 2.59 | 0.04 | 2.63 | 4151 |
| CGC | R | 0.09 | 3.69 | 87 | 0.12 | 0.06 | 2.42 | 102 | 0.08 | 0.21 | 10.60 | 1007 | 0.34 | 0.52 | 31.08 | 49083 |
| CGG | R | 0.04 | 1.61 | 38 | 0.16 | 0.04 | 1.40 | 59 | 0.14 | 0.13 | 6.49 | 617 | 0.65 | 0.17 | 9.91 | 15656 |
| CGT | R | 0.63 | 26.68 | 630 | 2.04 | 0.36 | 13.63 | 575 | 1.04 | 0.36 | 18.49 | 1757 | 1.41 | 0.22 | 13.08 | 20659 |
| AGC | S | 0.14 | 8.77 | 207 | 0.40 | 0.09 | 5.57 | 235 | 0.26 | 0.13 | 8.83 | 839 | 0.40 | 0.38 | 21.83 | 34478 |
| AGT | S | 0.19 | 11.48 | 271 | 2.62 | 0.25 | 16.31 | 688 | 3.72 | 0.21 | 14.03 | 1333 | 3.20 | 0.08 | 4.39 | 6929 |
| TCA | S | 0.17 | 10.55 | 249 | 2.63 | 0.22 | 14.06 | 593 | 3.51 | 0.14 | 9.05 | 860 | 2.26 | 0.07 | 4.01 | 6333 |
| TCC | S | 0.13 | 7.96 | 188 | 0.72 | 0.06 | 4.10 | 173 | 0.37 | 0.15 | 9.81 | 932 | 0.89 | 0.19 | 11.05 | 17449 |
| TCG | S | 0.04 | 2.33 | 55 | 0.20 | 0.03 | 1.71 | 72 | 0.15 | 0.15 | 9.93 | 944 | 0.86 | 0.20 | 11.56 | 18263 |

| Codon | AA[a] | ϕKp27 | | | | ϕKp34 | | | | ϕKp24 | | | | *K. pneumoniae* HS11286 | | |
|---|---|---|---|---|---|---|---|---|---|---|---|---|---|---|---|---|
| | | Fraction[b] | Frequency | No. | Ratio to bacteria | Fraction | Frequency | No. | Ratio to bacteria | Fraction | Frequency | No. | Ratio to bacteria | Fraction | Frequency | No. |
| TCT | S | 0.33 | 19.95 | 471 | 4.46 | 0.35 | 22.68 | 957 | 5.07 | 0.22 | 14.92 | 1418 | 3.34 | 0.08 | 4.47 | 7062 |
| ACA | T | 0.18 | 11.65 | 275 | 4.35 | 0.40 | 22.14 | 934 | 8.27 | 0.19 | 12.65 | 1202 | 4.73 | 0.05 | 2.68 | 4226 |
| ACC | T | 0.27 | 18.04 | 426 | 0.58 | 0.12 | 6.87 | 290 | 0.22 | 0.31 | 20.25 | 1925 | 0.65 | 0.61 | 31.30 | 49424 |
| ACG | T | 0.05 | 3.47 | 82 | 0.27 | 0.06 | 3.46 | 146 | 0.27 | 0.24 | 16.04 | 1524 | 1.24 | 0.25 | 12.94 | 20435 |
| ACT | T | 0.50 | 33.16 | 783 | 6.94 | 0.42 | 23.61 | 996 | 4.94 | 0.26 | 17.25 | 1639 | 3.61 | 0.09 | 4.78 | 7550 |
| GTA | V | 0.38 | 23.42 | 553 | 3.42 | 0.33 | 21.90 | 924 | 3.20 | 0.22 | 15.32 | 1456 | 2.24 | 0.10 | 6.85 | 10811 |
| GTC | V | 0.12 | 7.16 | 169 | 0.34 | 0.10 | 6.28 | 265 | 0.30 | 0.26 | 17.53 | 1666 | 0.83 | 0.30 | 21.08 | 33297 |
| GTG | V | 0.20 | 11.99 | 283 | 0.37 | 0.12 | 7.94 | 335 | 0.25 | 0.18 | 12.44 | 1182 | 0.39 | 0.46 | 32.30 | 51014 |
| GTT | V | 0.30 | 18.59 | 439 | 1.80 | 0.45 | 29.70 | 1253 | 2.87 | 0.34 | 23.25 | 2210 | 2.25 | 0.15 | 10.33 | 16318 |
| TGG | W | 1.00 | 12.75 | 301 | 0.80 | 1.00 | 14.58 | 615 | 0.91 | 1.00 | 13.54 | 1287 | 0.85 | 1.00 | 15.96 | 25204 |
| TAC | Y | 0.39 | 14.02 | 331 | 1.08 | 0.26 | 12.11 | 511 | 0.93 | 0.54 | 21.70 | 2062 | 1.67 | 0.48 | 13.03 | 20574 |
| TAT | Y | 0.61 | 21.81 | 515 | 1.54 | 0.74 | 33.99 | 1434 | 2.39 | 0.46 | 18.66 | 1773 | 1.31 | 0.52 | 14.21 | 22439 |
| TAA | *[c] | 0.70 | 2.75 | 65 | 1.39 | 0.59 | 3.46 | 146 | 1.75 | 0.66 | 2.57 | 244 | 1.30 | 0.52 | 1.97 | 3117 |
| TAG | * | 0.07 | 0.25 | 6 | 0.49 | 0.10 | 0.57 | 24 | 1.11 | 0.10 | 0.38 | 36 | 0.74 | 0.13 | 0.51 | 811 |
| TGA | * | 0.24 | 0.93 | 22 | 0.70 | 0.32 | 1.85 | 78 | 1.39 | 0.25 | 0.97 | 92 | 0.73 | 0.35 | 1.34 | 2108 |
| TAA | * | 0.70 | 2.75 | 65 | - | - | - | - | - | - | - | - | - | - | - | - |
| TAG | * | 0.07 | 0.25 | 6 | - | - | - | - | - | - | - | - | - | - | - | - |
| TGA | * | 0.24 | 0.93 | 22 | - | - | - | - | - | - | - | - | - | - | - | - |

[a] AA, amino acid

[b] Fraction, proportion of usage of the codon among the set of codons which code for this codon's amino acid; Frequency, number of specific codon divided by total number of codons in the sequence; No, total number of times the specific codon is observed in the sequence.

[c] *, stop codon

### References


1. Chaikeeratisak, V., Nguyen, K., Egan, M. E., Erb, M. L., Vavilina, A. and Pogliano, J. 2017, The phage nucleus and tubulin spindle are conserved among large pseudomonas phages. *Cell Rep.*, **20**, 1563-1571.
2. Malone, L. M., Warring, S. L., Jackson, S. A., et al. 2020, A jumbo phage that forms a nucleus-like structure evades CRISPR–Cas DNA targeting but is vulnerable to type III RNA-based immunity. *Nat. Microbiol.*, **5**, 48-55.
